# The auxin transporter PIN1 and the cytokinin transporter AZG1 interact to regulate the root stress response

**DOI:** 10.1101/2020.10.22.350363

**Authors:** TM Tessi, M Shahriari, VG Maurino, E Meissner, O Novak, T Pasternak, BS Schumacher, NS Flubacher, M Nautscher, A Williams, Z Kazimierczak, M Strnad, JO Thumfart, K Palme, M Desimone, WD Teale

**Author notes:** These authors contributed equally to this work (TM Tessi, M Shahriari, VG Maurino). Labormedizinisches Zentrum Ostschweiz, Lagerstrasse 30, 9470 Buchs SG, Switzerland.

## Abstract

Root system development is crucial for optimal growth and yield in plants, especially in sub-optimal soil conditions. The architecture of a root system is environmentally responsive, enabling the plant to exploit regions of high nutrient density whilst simultaneously minimizing abiotic stress. Despite the vital contribution of root systems to the growth of both model and crop species, we know little of the mechanisms which regulate their architecture. One factor which is relatively well understood is the transport of auxin, a plant growth regulator which defines the frequency of lateral root (LR) initiation and the rate of LR outgrowth. Here we describe a search for proteins which regulate RSA by interacting directly with a key auxin transporter, PIN1. The native separation of PIN1 identified several co-purifying proteins. Among them, AZG1 was subsequently confirmed as a PIN1 interactor. AZG1-GFP fusions co-localized with PIN1 in procambium cells of the root meristem. Roots of *azg1* plants contained less PIN1 and blocking proteolysis restored PIN1 levels, observations which are consistent with PIN1 being stabilized by AZG1 in the plasma membrane. Furthermore, we show that AZG1 is a cytokinin import protein; accordingly, *azg1* plants are insensitive to exogenously applied cytokinin. In wild-type plants, the frequency of LRs falls with increasing salt concentration, a response which is not observed in *azg1 x azg2* plants, although their drought response is unimpaired. This report therefore identifies a potential point for auxin:cytokinin crosstalk in the environmentally-responsive determination of root system architecture.

## Introduction

Soil salinity limits crop yields in many agricultural systems (Rozema and Flowers, 2008). In shaping root system architecture (RSA), salinity defines the surface area of the plant which is able to take nutrients and water from the soil. RSA is determined by complex signalling networks, guiding main and lateral roots to develop in response to NaCl concentration (Zolla et al., 2010; Julkowska and Testerink, 2017).

Factors which determine RSA, such as branching and growth rates are controlled by signals which are mediated by plant hormones. For example, artificially induced concentration maxima of the plant hormone auxin in root pericycle cells are necessary and sufficient to specify LR founder cells (Dubrovsky et al., 2008). Similar auxin maxima are generated naturally in protoxylem pole cells through an interaction between developmentally-defined auxin signaling oscillations in the root apical meristem (RAM) and the action of auxin transport proteins (Xuan et al., 2020). Throughout LR development, the generation and maintenance of dynamic auxin concentration maxima over several cell types is required for the correct initiation and development of a LR primordium (Benkova et al., 2003). The activity of several auxin transporters (particularly those of the PIN family) is necessary for these specific patterns of auxin distribution (Blilou et al., 2005; Benkova et al., 2003). A polar localization of PIN proteins in the plasma membrane directs the flux of auxin to define the site of auxin concentration maxima (Grieneisen et al, 2007; Petrasek et al, 2006). Furthermore, the distribution of several PIN proteins among polar plasma membrane domains is environmentally sensitive, acting to mediate RSA in response to gravity and mechanical obstacles in the soil (Ditengou et al., 2008; Friml et al., 2002; Ottenschläger et al., 2003). This sensitivity suggests a mechanism through which environmental factors may influence the rate at which roots grow and the extent to which they branch.

Auxin is counteracted by cytokinin during root development, with the ratio of the two hormones being a particularly important factor in the shaping of RSA (Aloni et al., 2006; dello Ioio et al., 2008; Muller & Sheen, 2008). Cytokinins repress LR initiation through the action of CRE1/AHK4, an endoplasmic reticulum-localized receptor (Caesar et al, 2011; Laplaze et al, 2007). The cross-talk between auxin and cytokinin signaling cascades is complex, with the pathways having a reciprocal influence (Muller and Sheen, 2008; dello Ioio et al., 2008). *PIN* transcription is regulated by cytokinin to regulate root development (Bishopp et al, 2011a). However, in addition to exerting transcriptional control over polar auxin transport, cytokinins also promote the degradation of PIN1 in regenerating root tissue (Marhavý et al., 2011), potentially regulating the polarity of PIN proteins in developing LRs (Marhavý et al., 2014). However, although PIN1 phosphorylation status determines sensitivity to cytokinin in this respect, the proteins which directly mediate cytokinin-mediated sub-cellular PIN1 localisation remain obscure (Marhavý et al., 2014). Indeed, over and above their links to cytokinin signaling, proteins which interact directly with PINs remain largely elusive.

The dedicated transport of cytokinin is presently largely characterized by the loading of *trans*-zeatin (tZ) into xylem cells in roots (Zhang et al., 2014; Ko et al., 2014), and the loading of N^6^(Δ^2^-isopentyl)adenine into phloem cells in leaves (Bürkle et al., 2003). Such long distance transport is thought to carry environmental information, such as on nutrient status, through the body of the plant. Short distance cytokinin transport has also been shown to play a role in the coordination of internal developmental cues; PURINE PERMEASE 14 (PUP14) is active in the developing embryo, and may even be involved in the relay of cytokinin signaling responses (Zürcher et al., 2016).

The generation of spatially distinct auxin and CK signalling domains is a key feature of vascular patterning (De Rybel et al., 2014; Ohashi-Ito et al., 2014), with the xylem axis being a region of high auxin signaling activity, and cells of the flanking cambial domains containing relatively high CK signaling activity (Bishopp et al., 2011a). Auxin-CK therefore appear to interact to drive distinct domains of root development, but a key question remains how these domains are coordinated both in the main root axis (De Rybel et al., 2014; Ohashi-Ito et al., 2014) and the LR (Chang et al., 2015, Bielach et al., 2012).

In this study we identify *Arabidopsis thaliana* AZG1 as a novel cytokinin import protein which directly interacts with PIN1. Our results are consistent with the hypothesis that AZG1 acts to stabilize PIN1 in the plasma membrane, potentially acting to sharpen the spatially separated boundary between auxin in protoxylem cells and cytokinin signaling maxima in procambial cells of the RAM. Moreover, we found that AZG1 is necessary to regulate LR density in response to NaCl, conferring sensitivity to salinity but not to drought.

## Materials and methods

### Preparation of Arabidopsis plasma membranes

Dark-grown MM2d *Arabidopsis thaliana* cell cultures (Menges and Murray, 2002) were harvested and washed with an equal volume of cold 20 mM KCl, 5 mM EDTA before being lysed at 2°C with an automatic cell pressure lysis machine at 0.3 KBar (Constant Systems) in homogenization buffer (50 mM Tris adjusted to pH 8.0 with MES, 500 mM sucrose, 10% glycerol, 20 mM EDTA, 50 mM NaF, 0.6% PVP (Mw 30,000 – 40,000), 10 mM ascorbic acid) containing a protease inhibitor cocktail (Roche). All subsequent steps were conducted on ice. Homogenate was centrifuged at 10,000 x g at 4°C for 10 minutes and filtered through two layers of Miracloth (Calbiochem) before centrifugation at 40,000 x g for one hour at 4°C. Pellets were resuspended in microsome buffer (5 mM phosphate phosphate buffer pH 7.8, containing 330 mM sucrose, 2mM DTT, 10 mM NaF) using 1ml of buffer for every 20g of cells lysed (fresh weight). Solubilized protein complexes were prepared by treating microsomes with 0.5% Triton X-114 under gentle rotation for one hour before centrifugation at 100,000 x g for 30 minutes. Plasma membranes were prepared by two-phase partitioning (Dextan T500/PEG3350) according to Kjellbom and Larsson (1984).

### Fractionation of Arabidopsis membrane protein complexes by free-flow electrophoresis

Isoelectric focussing (IEF)-free-flow electrophoresis (FFE) was conducted according to the manufacturer’s instructions (BD) in a chamber of 0.5 mm thickness. Laminar flow of buffer was buffer was verified with 0.1 % SPADNS and the chamber was treated with 0.05 % HPMC before IEF-FFE amphylites (pH 3-9; BD) were applied to inlets 2-6 of the chamber at a flow rate of 60 ml per hour. Protein complexes were kept soluble by the addition of 0.8 % n-octyl-ẞ-D-pyranoside to all amphylites. A potential difference of 600 V was applied across the chamber and the pH gradient was verified using pH indicator dye mixture (BD). For each experiment, 300 µl of microsomal sample was introduced to the chamber with a peristaltic pump at a flow rate of 700 µl per hour (between 1 and 2 mg protein per hour). Separated protein complexes were collected in a 96-well plate and each well was precipitated by the addition of a 10% volume of saturated trichloroacetic acid (*aq*) and a 1% volume of 2% sodium deoxycholate before incubation on ice for one hour and centrifugation at 20,000 x g for 30 min. Samples were visualized by SDS-PAGE and subsequent silver staining. PIN1-containing fractions were identified by western blotting with an anti-PIN1 monoclonal antibody as previously described (Blilou *et al*., 2005), and submitted for MS/MS analysis along with the nearest fraction of a higher protein concentration in which no PIN1 signal was detected.

### Fractionation of Arabidopsis membrane protein complexes by blue-native PAGE electrophoresis

Microsomal pellets were resuspended in ice-cold 20 mM bis-Tris pH 7.0 containing 500 µM ɛ-aminocaproic acid, 20 mM NaCl, 2 mM EDTA and 10% glycerol and 0.6% dodecyl maltoside before centrifugation at 100,000 x g for 30 min. Samples were diluted 1:1 with 750 µM aminocaproic acid containing 5% (w/v) coomassie G250. First dimension BN-PAGE proceeded according to Swamy *et al*. (2006). After separation, lanes were longitudinally cut into two strips. One strip was subjected to SDS-PAGE and western blotting as described above in order to identify regions which contained PIN1, the other was used for MS/MS. Corresponding strips, of a higher protein concentration in which no PIN1 signal was detected, were also submitted for MS/MS analysis.

### Mass spectrometry

The respective gel parts were in-gel digested with trypsin. Resulting peptides were extracted from the gel parts and separated via HPLC (Shevchenko et al., 1996; Blagoev et al., 2004). For this analyses an UltiMate3000 HPLC system with Chromeleon software (Dionex, Germany) was used. Peptides were eluted with 300 nl/min flow rate and a gradient from 3 % to 30 % buffer A (80 % (v/v) acetonitrile with 0.5 % acetic acid) in 60 minutes on a self-packed C18 column (Reprosil material, 3 μm particle size, packed in a fused silicate column with 75 μm inner diameter, approximately 15 cm length and a nanospray emitter tip with an 8 μm opening, New Objective, USA). Buffer A was water containing 0.5 % (v/v) acetic acid. The eluting peptides were transferred to a high-resolution mass spectrometer via a nano-elctrospray ion source with 5 kV ioniziation current. Mass spectrometry was performed in positive ion mode on a LTQ-Orbitrap hybrid mass spectrometer (ThermoFisher Scientific, Germany) using the manufacturer’s software. Peptides are identified via their exact masses and fragment ion patterns from tandem mass spectrometry fragment spectra (IDA mode). MASCOT server (Matrixscience Inc., UK) with the ncbi nr database, restricted to preen plants, was used for protein identification (Perkins et al., 1999).

### Plant lines

The experiments were performed with the wild-type Arabidopsis (WT) ecotype Columbia 0 (*Col-0*) and the *azg1* knockout t-DNA insertion lines *azg1-1* (SAIL_114E03; Tessi et al., 2020), *azg1-2* (GK-681A06; *Tessi et al., 2020)*. Unless explicitly indicated, experiments used *azg1-1*. A double t-DNA insertion line *azg1-1 x azg2-1*, obtained by genetic crossing of *azg1-1* and *azg2-1* (SALK 000904; Tessi et al., 2020), was included in the analysis. For localization experiments the GUS lines *ARR5pro:GUS*, *AZG1pro:GUS* and *azg1.1xARR5pro:GUS* were used. *TCSnpro:GFP* was kindly provided by Dr. Müller (Zürcher, 2013).

### Generation of transgenic plants overexpressing AZG1

AZG1 coding region was amplified from Arabidopsis root cDNA using Platinum Pfx DNA polymerase (Invitrogen) and the following primer combination: AZG1SmaI-F (5′-ATCCCGGGATGGAGCAACAGCAACAACAACAA-3) and AtAzg2XbaI-R (5′-AGTCTAGACTAAACGGTAGTATCAATCTCACTA-3). The primers introduce the unique restriction sites SmaI and XbaI at the 5′ and 3′end, respectively. The amplified product was cloned into pCR-Blunt II-TOPO (Invitrogen) and sequenced using the PRISM fluorescent dye-terminator system (Applied Biosystems). The 1.4 kb fragment was subcloned into a modified version of the binary vector pGreen II (Fahnenstich et al., 2007) bearing the BASTA resistance gene, using the SmaI and XbaI sites between the CaMV35S promoter and the Octopin-Synthetase-Terminator from the pBinAR vector. The resulting modified version of the pGreenII vector was called pGreenII-35S-nosBAR and the plasmid containing *AZG1* was called *p35S:AZG1*. The plasmid *p35S:AZG1* was introduced into Arabidopsis by *Agrobacterium tumefaciens* (GV3101) mediated transformation, using the vacuum infiltration method. Transformants were selected for resistance to BASTA. DNA was extracted from leaf material collected from selected plants and used for PCR analyses. Plants containing the transgene were transplanted and allowed to self. Seeds from the primary (T1) generation were sown and resultant T2 plants were subjected to another round of BASTA selection and characterization by means of PCR. The process was repeated to obtain non-segregating T3 transgenic lines. All further analyses were performed with homozygous T3 transgene plants.

### Generation of reporter lines

A DNA fragment containing a 1713 bp upstream and 179 bp downstream of the AZG1 start codon was amplified by RT-PCR with the primers AZG1-GUS_f 5′-CACCACACTGCGGCCTAAGAGAACAAACT-3′ and AZG1-GUS_rev 5′-GGTACCGCCACTTTCTTAACCATG-3′. This fragment was cloned into pGWB3, a gateway-compatible binary vector designed for promoter-driven expression of the GUS gene (kindly provided by T. Nakagawa, Shimane University, Izumo, Japan). Cloning using gateway vectors was done using reagents and protocols from Invitrogen (Carlsbad, California, United States). The plasmid containing the chimeric *AZG1pro:GUS* gene was introduced into *A. thaliana* by *Agrobacterium tumefaciens* (GV3101) mediated transformation using the vacuum infiltration method. Transgenic lines were selected with Kanamycin and analysed for GUS activity.

To study the subcellular localization the *U10pro:AZG1-GF*P was generated. The *AZG1* full coding sequence was cloned using Gateway cloning into pDONR207 entry vector and subcloned into pUBC-GW-GFP-DEST (Grefen et al., 2010). To drive the expression of *AZG-GFP* or *AZG1-YFP* with its native promoter, 1,5 kb of *AZG1* promoter and the full coding sequence of an AZG-GFP or AZG1-YFP translational fusion were cloned in a single amplicon and introduced in pDONR207. The resulting *AZG1pro:AZG1-YFP* or *AZG1pro:AZG1-YFP* fusion was subcloned into the pMDC107 destination vector. All constructs were verified by sequencing.

### Plant growth conditions

Plants were grown from seeds sown on Arabidopsis medium (AM) (2.2 g MS salts per litre, 2.5 mM MES pH 5.7, 15 mM sucrose containing 1.3% agar). After surface sterilization, seeds were incubated in dark overnight at 10°C. All plates were then transferred into vertical position in a growth chamber (Versatile Environmental Test Chamber, Sanyo, Japan) with long-day conditions of 16 h light and 8 h dark at 22°C.

Western blotting of PIN1 in plants proceeded according to Blilou et al. (2005). Two-day old seedlings were homogenised directly in Laemmli buffer (100 µl per 10 mg), centrifuged for 10 minutes at 24000 x g and loaded directly onto an SDS-PAGE gel. MG132 at 20 µM and cyclohexamide (50 µM) treatments were for 90 minutes immediately prior to homogenization.

### Root system phenotyping

To study the root system phenotype plant were grown vertically in Petri dishes. For general phenotyping under standard conditions MS 0,5 without Nitrogen + 20 mM KNO_3_ media was used. Plant were grown in 8-hour light 16-hour dark regime for 10 days after germination (dag). To study the effect of CK, plants were grown for 12 dag in 0.5x MS with or without the addition of 200 nM tZ.

For the study of root system reorganization, seedlings were grown in Petri dishes for 7 dag before the ablation of the distal 5 mm of the main root. Seedling were photographed 10 days after ablation. The same conditions were used to analyse the response of *TCSnpro:GFP* and *AZG1pro:GUS* after ablation. Root systems were measured and analyzed using ImageJ-FIJI.

### Immunolocalization

Four-day old Arabidopsis seedlings were fixed twice under vacuum for 2 minutes in MTSB-T (1x MTSB, 4 % formaldehyde, 1 % Triton-X-100) followed by 40 minutes’ incubation in the dark at room temperature. Afterwards all samples were washed with water and with MTSB-T and were transferred to an InsituPro pipetting robot (Intavis AG, Germany) with 650 µl buffer per well. The following treatments were then performed: 5 washes for 10 minutes in MTSB, 40 minutes’ digestion in digestion solution (10.5 mg macerozyme R-10 and 10.5 mg driselase in 1 ml H_2_O) were centrifuged for 4 minutes at 1000 rpm. The supernatant was added to 1 ml 5 mM MES and made up to 7 ml with H_2_O), 5 washes for 12 minutes in MTSB, 2 incubations for 25 minutes in 2x MTSB, containing 20% DMSO and 0.6% NP-40, five washes for 10 minutes in MTSB, 1 hour incubation in MTSB containing 2 % BSA, 5 hours incubation in MTSB containing 2 % BSA containing either an anti-GFP (Roche) or an anti-PIN1 monoclonal antibody at 0.1 %, 6 washes for 12 minutes in MTSB, 4 hour incubation in MTSB containing 2 % BSA and 0.5 % Alexa-conjugated anti-mouse secondary antibody (Invitrogen), 9 washes for 15 minutes in MTSB, 7 washes for 15 minutes in deionized water. Coimmunolocalization was performed with a rabbit anti GFP primary antibody (Roche) and appropriate combination of Alexa-conjugated secondary antibodies (Invitrogen).

### Callus generation

Root segments of 5 mm were excised and incubated on AM medium containing 2,3 µM 2,4-dichlorophenoxyacetic acid and 230 nM kinetin. The resulting calli were propagated by placing freshly cut sections on new media plates every three weeks.

### GUS staining

Plants were incubated for 20 minutes in 90 % acetone (*aq*) at room temperature before infiltration under vacuum for 30 min in 50 mM NaPO_4_ (pH 7,2), 2 mM Potassium-Ferrocyanide, 2 mM Potassium-Ferricyanide, 0,2 % Triton-X-100, 2 mM X-Gluc. After incubation for 3 h in the dark plants were washed and sequentially incubated for 30 minutes in 70 %, 80%, 90% and 100% ethanol. The stained plants were stored in 100 % ethanol before mounting on glass slides with the SlowFade Antifade Kit (life technologies, USA). All GUS staining was observed and recorded with an AXIO Imager.A1 (Zeiss, Germany) using a Zeiss EC Plan-NEOFLUAR 10x/0,3 objective and Zeiss PLAN APOCHROMAT 20x/0.8 objective with use of Zeiss Zen 2 software. All pictures were taken with a Zeiss Axiocam 105 colour Camera.

### iRoCS

Propidium iodide-mediated staining of cell boundaries was performed in order to record 3D images for iRoCS analysis (Schmidt et al. 2014). Root tips of 1cm were transferred to propidium iodide fixative solution, vacuum infiltration was applied twice for 2 minutes and roots were incubated for at least 12 h at 4°C. Samples were sequentially washed in 80 %, 50 % and 30 % ethanol, each for 5 minutes. Roots were then rinsed twice in water for 5 minutes and incubated in 20 mM KPO4 buffer (pH 7.0), 2 mM NaCl, 0.25 mM CaCl2 and 0.1 mg/ml amylase overnight at 37°C to remove starch from the root tip. The roots were rinsed twice in water for 5 minutes and then acidified by the addition of periodic acid to a concentration of 1% for 30 minutes at room temperature. Roots were again washed twice in water for 5 minutes and moved to Schiff reagent containing 0.1 mM propidium iodide for 30 minutes. From this stage, roots were protected from light. The samples were rinsed twice in water for 5 minutes and then incubated in aqueous 73% (^w^/_v_) chloral hydrate for 2 h. Glycerol was added to 20 % and roots were incubated overnight before mounting in in chloral hydrate solution containing 20 % glycerol.

Propidium iodide stained roots were detected by a LSM 510 META laser scanning microscope (Zeiss, Germany) and a 40x/1.3 DIC objective with oil immersion. Stacks of images were reconstructed into a three dimensional image as previously described (Schmidt et al., 2014) and automatically stitched by ZEN 2010 (Carl Zeiss MircroImaging GmbH).

Analysis of propidium iodide stained roots was performed by the intrinsic root coordinate system (iRoCS) according to Schmidt *et al.* (2014). After manual marking of the quiescent centre, the iRoCS Toolbox automatically detected cells and attached them to different cell layers in the root (epidermis with atrichoblasts and trichoblasts, cortex, endodermis, pericycle and vascular tissue) or designated them under- or over segmented. The marking by iRoCS was then manually corrected and under- as well as over-segmented cells were manually excluded from subsequent analysis. Data of cell parameters were analyzed with R-Studio.

### Cytokinin extraction, purification and quantitative analysis by UPLC-MS/MS

Wild-type and *azg1-1* plants were grown on MS plates or alternatively MS supplemented with 200 nM tZ. Four independent biological samples made up of 1 g pooled rosette leaves were harvested after 26 days and kept frozen at −80°C. The procedure used for cytokinin analysis was a modification of the method described by Faiss et al. (1997). Freeze–dried plant material was homogenized in liquid nitrogen and extracted in ice-cold 70% (^v^/_v_) ethanol. Deuterium-labelled CK internal standards (Olchemim Ltd., Czech Republic) were added, each at 5 pmol per sample to check the recovery during purification and to validate the determination. The standards were [2H5]tZ, [2H5]tZR, [2H5]tZ9G, [^2^H5]tZOG, [^2^H5]tZROG, [^2^H5]tZRMP, [^2^H3]DHZ, [^2^H3]DHZR, [^2^H3]DHZ9G, [^2^H3]DHZOG, [^2^H3]DHZROG, [^2^H3]DHZRMP, [^2^H6]iP, [^2^H6]iPR, [^2^H6]iP9G, [^2^H6]iPRMP, [^2^H7]BA, [^2^H7]BAR,[^2^H7]BA9G, [^2^H7]BARMP, [^15^N4]mT, and [^15^N4]oT. All topolins were analysed using internal deuterium standards for [^15^N4]mT and [^15^N4]oT as no other labelled standards were available. Therefore, the values of other topolin metabolites may have an error load which originates from imperfect internal standardization. After 3 h extraction, the homogenate was centrifuged (15,000 g at 4 °C) and the pellets were re-extracted. The combined supernatants were concentrated to approximately 1.0 ml by rotary evaporation under vacuum at 35 °C. The samples were diluted to 20 ml with ammonium acetate buffer (40 mM, pH 6.5). The extracts were purified using a combined (diethylamino)ethyl (DEAE)-Sephadex (Sigma-Aldrich, St. Louis, MA, USA) (1.0 x 5.0 cm)-octadecylsilica (0.5 x 1.5 cm) column and immunoaffinity chromatography (IAC) based on wide-range specific monoclonal antibodies against cytokinins (Faiss et al., 1997). This resulted in 3 fractions: (1) the free bases and 9-glycosides (fraction B), (2) a nucleotide fraction (NT) and (3) an O-glucoside fraction (OG). The metabolic eluates from the IAC columns were evaporated to dryness and dissolved in 20 μl of the mobile phase used for quantitative analysis.

The cytokinin fractions were analysed by ultra-performance liquid chromatography (UPLC) (ACQUITY UPLCTM; Waters, Milford, MA, USA) linked to a Quattro microTM API (Waters, Milford, MA, USA) triple quadrupole mass spectrometer equipped with an electrospray interface. The purified samples were dissolved in 15 μl MeOH/H_2_O (30/70) and 10 μl of each sample was injected onto a C18 reversed-phase column (Acquity UPLCTM is based on a combination of high pressure and small bridged ethylsiloxane/silica hybrid particles; BEH Shield RP18; 1.7 μm; 2.1 х 150 mm; Waters). The column was eluted with a linear gradient of 15mM ammonium formate (pH 4.0, A) and methanol (B), with retention times for the monitored compounds ranging from 2.50 to 6.50 min. The binary gradient (0 min, 10 % B; 0-8 min, 50 % B) was applied with a flow-rate of 0.25 ml/min and a column temperature of 40 °C. Quantification was obtained by multiple reaction monitoring of [M+H]+ and the appropriate product ion. For selective MRM experiments, optimal conditions were as follows: capillary voltage 0.6 kV, source/desolvation gas temperature 100/350 °C, cone/desolvation gas 2.0/550 L/h, LM/HM resolution 12.5, ion energy 1 0.3 V, ion energy 2 1.5 V, entrance 2.0 V, exit 2.0 V, multiplier 650 eV. The dwell time, cone voltage, and collision energy in collision cell corresponding to exact diagnostic transition were optimized for each cytokinin. On the basis of retention time stability, the chromatographic run was split into eight retention windows. The dwell time of each MRM channel has been calculated to obtain 16 scan points per peak during which time the inter channel delay was 0.1 s. In MRM mode, the limit of detection for most of cytokinins was below 5.0 fmol and the linear range was at least five orders of magnitude.

## Results

### AZG1 interacts directly with PIN1

PIN1-mediated polar auxin transport is an important factor which determines the size and shape of a root system. Once positioned correctly, auxin is likely to interact with a dense lattice of morphogenic signaling cascades. Prominent among these interacting signaling cascades are those stimulated by cytokinin, which affect auxin signaling and feed back onto auxin transport both transcriptionally and post-translationally (dello Ioio et al., 2008; Marhavý et al., 2014). In order to search for direct post-translational regulators of polar auxin transport, we searched for proteins which interacted with PIN1 in plasma membrane extracts from dark-grown Arabidopsis cell cultures. These membrane preparations were solubilized with 0.5 % Triton X-114; conditions which preserve the integrity of PIN1-containing membrane complexes. After the removal of detergent-insoluble membranes by centrifugation, the remaining soluble protein complexes were separated either according to their size (by blue-native PAGE) or their net charge (by isoelectrofocusing (IEF)-free flow electrophoresis (FFE)). FFE was chosen over capillary electrophoresis as it was able to maintain membrane protein solubility as complexes approached their isoelectric point. Fractions containing PIN1 were identified by western blotting and corresponded to a molecular mass of approximately 440 KDa and a pI of 5.4 (Figure 1, A and B). Proteins of PIN1-containing fractions were identified by MS/MS spectrometry, and compared with the nearest fraction which did not contain PIN1. Proteins were considered to be putative PIN1 interactors if i) they were identified in both BN-PAGE and IEF-FFE experiments, ii) they were represented by more than one peptide in each experiment, and iii) were never identified in a control fraction. In total, peptides derived from 141 different proteins fulfilled these criteria after BN-PAGE fractionation, and from 133 proteins after IEF-FFE fractionation. The proteins which were found in both groups were classified as potential PIN1 interactors, of which there were 33 (Figure 1A and B). This report focuses on the physiological characterization of one of these proteins: a homolog of AzgA, a fungal purine transporter of the AZA-GUANINE RESISTANCE transporter family, named AZG1 (Mansfield et al., 2009; Tessi et al. 2020). AZG1 was identified by our MS/MS analysis with a total of seven peptides (Figure 1C).

**Figure 1.**
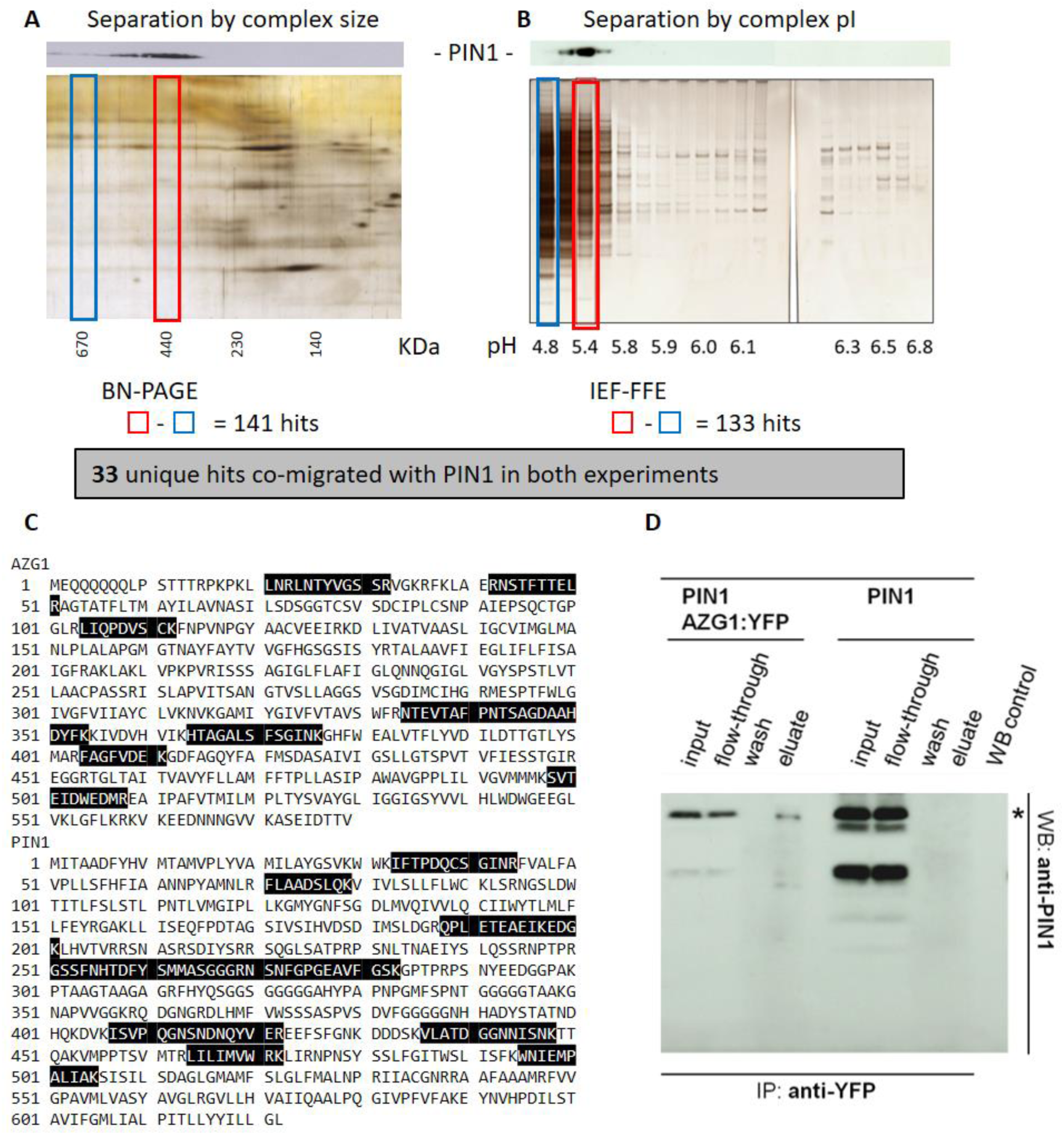
The identification of AZG1 as an interactor of PIN1. Arabidopsis microsomal protein comlexes were solubilized and separated by either (A) BN-PAGE or (B) IEF-FFE. Western blots used an anti-PIN1 primary antibody, total proteins (A and B, below) visualized with silver staining. Numbers of protein hits which were identified exclusively in PIN1-positive fractions (red boxes) and not in PIN1 negative fractions (blue boxes) for each method are given. C) Primary sequences of AZG1 and PIN1 indicating peptides identified in either BN-PAGE or IEF-FFE. D) Affinity purification of PIN1 using an anti-GFP antibody is dependent on the presence of a translational AZG1:YFP fusion. Proteins were extracted from tobacco mesophyll protoplasts transiently transformed with Arabidopsis coding sequences under the control of a 35S promoter. Asterisk indicates PIN1 signal. WB control shows untransformed cells.

In order to rule out an accidental co-migration of PIN1 and AZG1-containing protein complexes in both separation techniques, the proximity of the PIN1-AZG1 interaction was tested in an affinity-purification assay. Both PIN1-HA and a C-terminal AZG1-YFP fusion protein were transiently expressed in tobacco leaves, before microsomal proteins were solubilized with 1% dodecyl maltoside and AZG1-YFP was pulled-down with an immobilized anti-GFP antibody. Presence of PIN1-HA in the eluate, as determined by western blotting with an HRP-conjugated anti-HA antibody, was dependent on the presence of AZG1-YFP (Figure 1D), indicating a binary interaction. Not all of the 33 proteins which co-migrated with PIN1 in IEF-FFE and BN-PAGE were pulled down by PIN1 in this transient expression system. For example, two proteins with plausible functional links to auxin transport did not directly bind PIN1: SMT2, a sterol methyltransferase involved in cotyledon vein patterning (Carland et al., 2002) and DRP1A, a dynamin-like protein which has previously been shown to co-immunoprecipitate with PIN1, but is not thought to be a direct interactor (Mravec *et al*., 2011; Figure S1).

### AZG1 is a high affinity adenine importer, and CKs are strong competitors

To better understand any potential physiological relevance of the AZG1-PIN1 interaction, we characterized the transport activity of AZG1. The AZG family has been primarily described in the filamentous Ascomycete *Emericella nidulans*. The gene family take its name from the capability of EnAZGA to transport not only purines (adenine, guanine and hypoxanthine) but also their toxic analogs aza-guanine and aza-adenine (Cecchetto, 2004). In plants, AZG1 has also previously been described as a cellular purine importer (Mansfield et al., 2009).

In order to further characterize the capacity of AZG1 to import purine in to the cell, we set up a heterologous model of adenine uptake in yeast. Cells expressing AZG1 displayed a linear accumulation of radiolabeled adenine for the first three minutes of incubation; the accumulation rate was more than 10-fold higher than untransformed control cells, consistent with the results reported by Mansfield et al. (2009) (Figure 2A). The rate of adenine import was saturable by increasing substrate concentration. The deduced Michaelis-Menten kinetic parameters indicate a *K*_*m*_ of 1.62 ± 0.21 μM and a *V*_*max*_ of 16.2 ± 0.28 pmol/10^6^ cells/minute (Figure 2B). We therefore conclude that AZG1 is a high-affinity adenine transporter. To our knowledge, the *K*_*m*_ of AZG1 for adenine makes it the highest affinity purine transporter reported to date.

**Figure 2.**
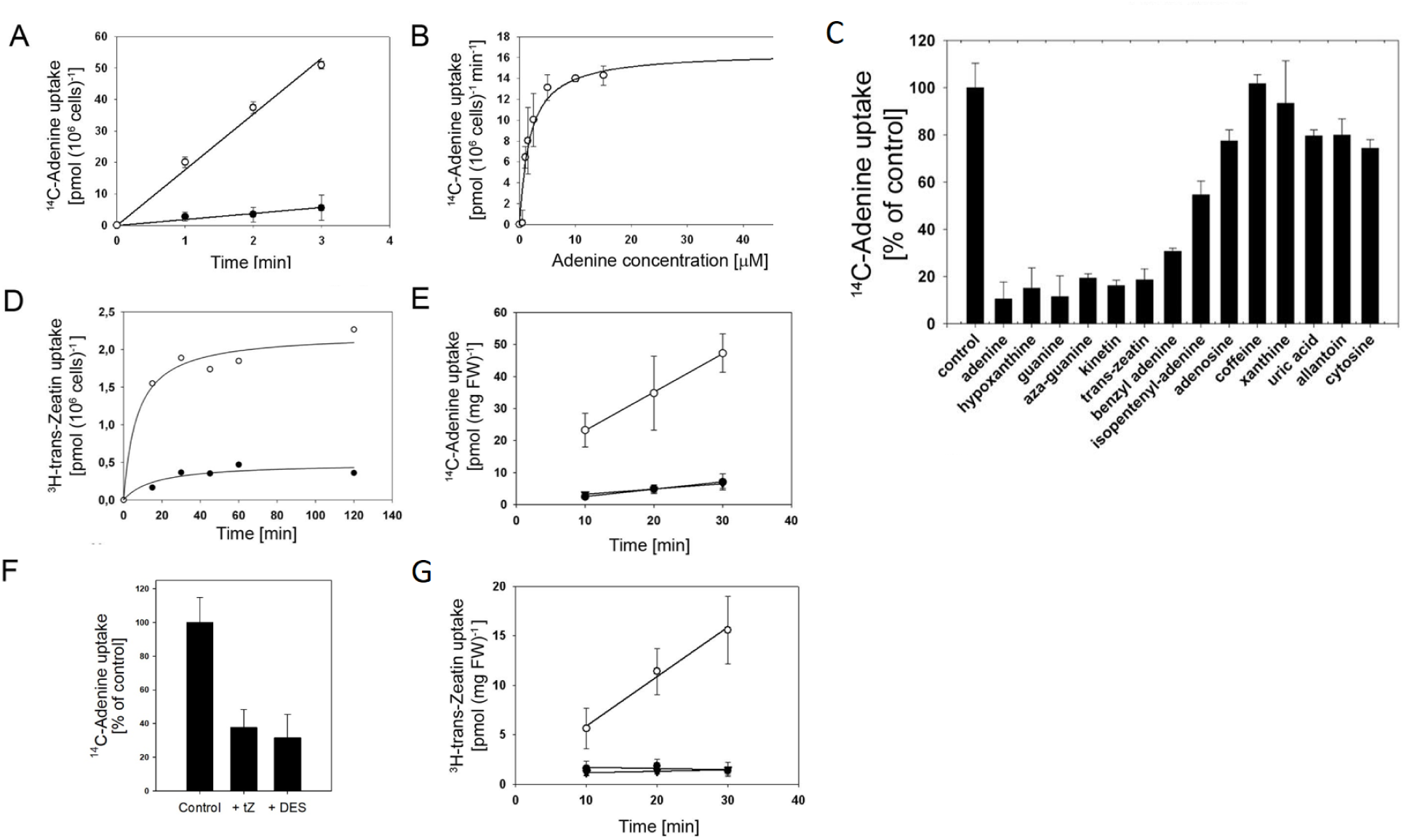
A) Yeast cells transformed with pESC-AZG1 (open circles) or with the empty vector (closed circles) were assayed for ^14^C-adenine uptake at 20 μM substrate concentration and pH 4. B) Concentration-dependent AZG1-mediated adenine uptake. Rates were calculated after the subtraction of background uptake rates (empty vector). C) AZG1-mediated uptake of ^14^C-adenine (20 μM) was determined in the presence of 10-fold excess (200 μM) of unlabelled compounds. Stated values have been normalized to uptake rates in the absence of competitors. D) Yeast cells transformed with pESC-AZG1 (open circles) or with the empty vector (closed circles) were assayed for ^3^H-trans-zeatin (tZ) uptake at 20 μM substrate concentration. (E) Uptake of ^14^C-adenine (E) or ^3^H-tZ. F) Proton dependence of ^14^C-adenine uptake and competition with tZ. ^14^C-Adenine (25 µM) net uptake by 35Spro:AZG1 plants as a percentage of wild type control. Also given are values in the presence of either diethylstilbestrol (DES) or 250 µM tZ. (G) into 14-day-old Arabidopsis seedlings. WT (closed circles), *azg1* (triangles) and 35S:AZG1-1 (open circles) seedlings were incubated with 20 μM ^14^C-adenine for the indicated time All experiments were run for 30 minutes. Values given are averages of four experiments, bars indicate standard error. Values represent the mean ± SD of three independent experiments.

The substrate specificity of AZG1 was investigated by determining the ability of a range of compounds to inhibit AZG1-mediated ^14^C-adenine uptake in yeast cells when present in a 10-fold excess (Figure 2C). Adenine, guanine and hypoxanthine were all strong competitors. Less effective were cytosine (a substrate for plant purine transporters, PUPs; Burkle et al., 2003; Zürcher et al., 2016), caffeine (which is also transported by PUPs), adenosine (a substrate of equilibrative nucleoside transporters, ENTs; Möhlmann et al., 2001), as well as xanthine, uric acid, uracil and allantoin (substrates for NATs; Maurino, et al., 2006; Niopek-Witz et al., 2014, and Ureide Permeases, UPS; Desimone et al., 2002); all only had a marginal effect on the transport of ^14^C-adenine. 8-Azaguanine strongly inhibited the AZG1-mediated cellular import of adenine, consistent with the resistance phenotype of overexpression and loss-of-function lines (Mansfield et al., 2009; Tessi et al., 2020). In general, the range of compounds which inhibited AZG1-mediated adenine accumulation was similar to that of EnAZGA (Cecchetto et al., 2004) suggesting a conserved protein function for eukaryotic AZG transporters.

### AZG1 is a cytokinin importer

Cytokinins form a class of plant growth regulators which share structural similarities with purines. Remarkably, aside from other purines, different cytokinin where the strongest inhibitors of AZG1-mediated cellular adenine import. In this respect, tZ and kinetin were highly effective competitors, followed by benzyl-adenine and pentenyl-adenine (Figure 2C). These data indicated that AZG1 most likely use these cytokinins as substrates. Therefore, we next measured the capacity of AZG1 to transport radiolabeled tZ. In this case, uptake of the maximal amount of label was reached after a few seconds, at which point a significant difference in the amount of accumulated radioactive tZ was observed between control and AZG1-expressing cells (Figure 2D).

To determine whether AZG1 was able to transport physiologically relevant purines *in planta*, 14-day-old seedlings were incubated with ^14^C-adenine. Both WT and *azg1-1* loss-of-function seedlings showed a linear uptake over 30 minutes, but no significant differences in the uptake rates could be observed (Figure 2E). It is possible that the accumulation of ^14^C-adenine in WT and loss-of-function seedlings was primarily mediated by proteins other than AZG1. In contrast, the transport rate of ^14^C-adenine in *35Spro:AZG1* seedlings was at least five times higher than that which was measured in WT plants (Figure 2E).

To test whether AZG1 was able to transport CK *in planta*, the yeast competitive uptake experiment was replicated in seedlings. Here, the transport of ^14^C-adenine in *35Spro:AZG1* seedlings was strongly inhibited by the addition of a 10-fold excess of unlabeled tZ, or after disruption of proton gradients by the addition of diethylstilbestrol (Figure 2F). These data suggest that tZ can compete with adenine for AZG1-dependent transport in seedlings.

To test whether cytokinins were being taken up into the cytosol, or were merely competing with adenine for membrane binding sites and not being transported, the capacity of AZG1 for cytokinin transport was addressed by measuring the uptake of ^3^H-tZ in 14-day-old seedlings. As was seen in the transport studies with ^14^C-adenine, WT and loss-of-function seedlings showed no significant differences in uptake, but AZG1 overexpressors exhibited an enhanced uptake capacity (Figure 2G). The uptake rate of ^3^H-tZ by AZG1 overexpressors was directly proportional to incubation time for at least 30 minutes and its accumulation rate was approximately six times higher than was measured in WT plants. These results demonstrate that AZG1 is able to drive cellular tZ uptake *in planta*.

We next tested whether the loss of AZG1 function altered the profile of cytokinin metabolites in the plant after exogenous treatment with tZ. Three-week old WT and AZG1 plants were grown in the presence of 200 nM tZ before rosette leaves were harvested and their cytokinin content measured. Consistent with the hypothesis that AZG1 functions as a cytokinin uptake protein in plants we observed a decrease in all tZ derivatives measured in the AZG1 plants when compared to the WT (Figure S2).

### AZG1 co-localizes with PIN1 in the plasma membrane of Arabidopsis root cells

PIN1 is polarly localized to the plasma membrane, from where it is thought continuously to recycle to endosomal compartments. In the root apical meristem (RAM), PIN1 is localized to the rootward face of stele cells, and is especially prominent in protoxylem pole cells. In order to ascertain whether AZG1 shares an expression domain with PIN1, plants were constructed in which the expression of AZG1-YFP was driven by its native promoter. When localized with an anti-GFP polyclonal antibody, AZG1-YFP was seen to localize to LR cap cells, epidermal cells and stele cells of the RAM, but was seen most distinctly in protophloem cells (Figure 3 A and B). AZG1 was also continuously expressed from the heart stage of the developing embryonic root, where its expression domain became gradually more refined through the torpedo stage to vascular cells in the mature embryonic seedling (Figure 3 C-E). Visualization of AZG1 after transient expression of in Arabidopsis protoplasts and roots with a *pU10:AZG1-GFP* construct indicated it also resided in the plasma membrane (Figure 3 F-K).

**Figure 3.**
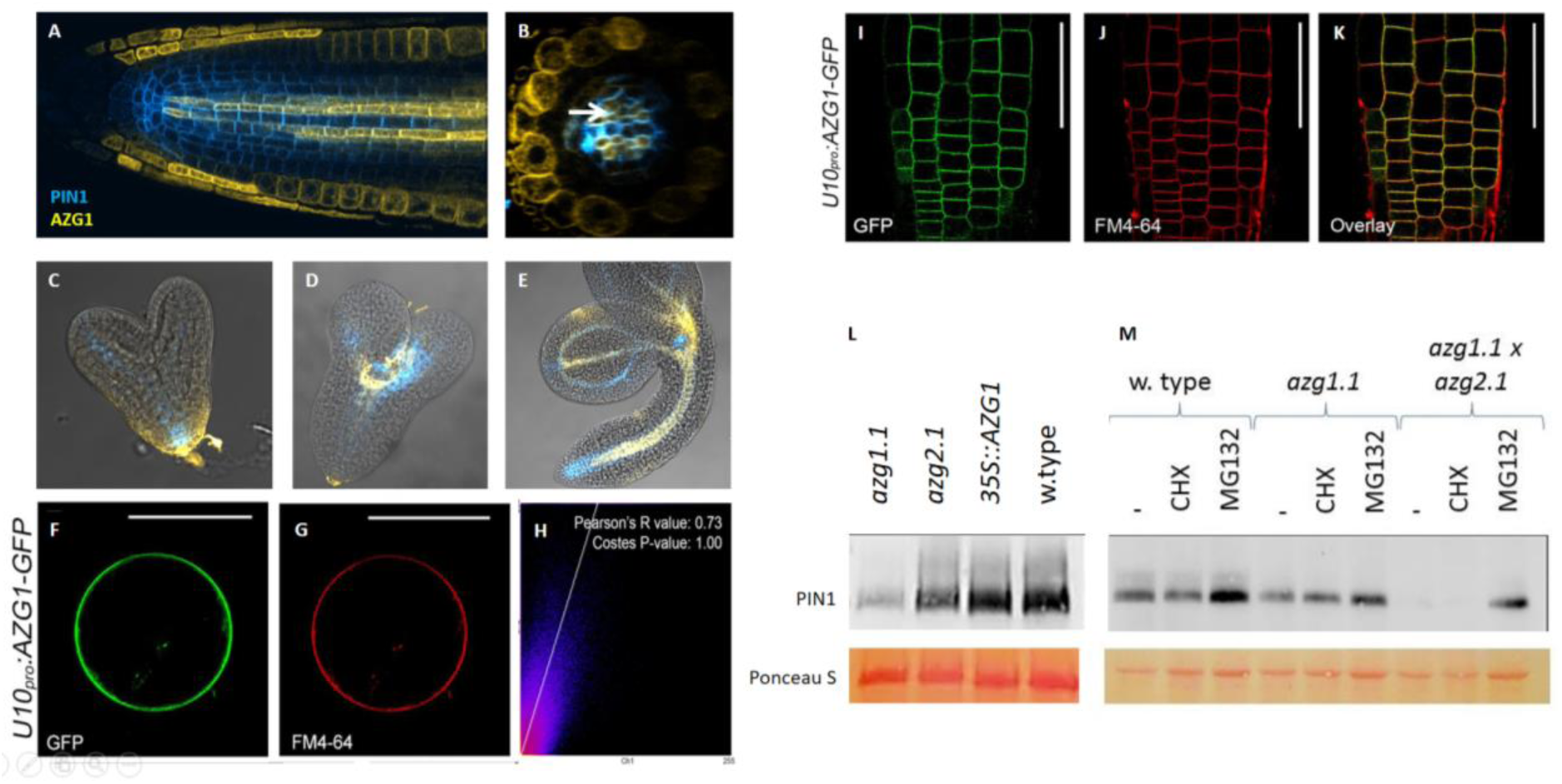
AZG1 *stabilizes PIN1 in co-expression domains*. Coimmunolocalization of AZG1:GFP (yellow) and PIN1 (blue) in A) the root apical meristem, B) the root apical meristem in radial section, Embryos at C) heart stage, D) torpedo stage, and E) mature stage. Subcellular localization of AZG1. F) Transient expression of *pU10:AZG1-GFP* construction in Arabidopsis root protoplasts compared to G) FM4-64 staining. H) Co-localization analysis of AZG1-GFP and FM4-64 marker of figures I and J performed using Coloc2 (FIJI). I) AZG1-GFP fusion under the control of the U10 promoter in epidermis cells of the meristematic zone compared to J) FM4-64 staining, and K) with the signals overlayed. Scale bars represent (F and G) 10 µm and (I-K) 50 µm. L) The relative amounts of PIN1 in 2-day old Arabidopsis seedlings. M) The relative amounts of PIN1 in 2-day old Arabidopsis seedlings after treatment with inhibitors of protein translation (CHX) and degradation (MG132). Loading control shows PVDF membranes stains with Ponceau S. Western blots were performed with an anti-PIN1 monoclonal antibody.

RT-PCR analysis revealed that *AZG1* was expressed in all tissues tested, with flowers and roots showing the highest amounts of transcript (Figure S3A). Expression of *pAZG1:GUS* was also observed in most tissues (Figure 4); primary root and LR apical meristems displayed distinct and different *pAZG1:GUS* staining patterns (Figure 4I-L). In seedlings the GUS staining broadened after 4 hours treatment with 200 nM tZ (Figure 4M and N). In the primary RAM, GUS staining was relatively evenly distributed, but the lateral RAM showed a narrower distribution of staining which was focused around the quiescent center (QC). In both types of root, CK induced *pAZG1*-dependent gene expression. A 90° rotation of plates induced *pAZG*-dependent gene expression in peripheral cells of the apical meristem of LRs (Figure 4O and 4P), but the gravitropic response of *AZG1* loss of function plants was only mildly affected (Figure S3B).

**Figure 4.**
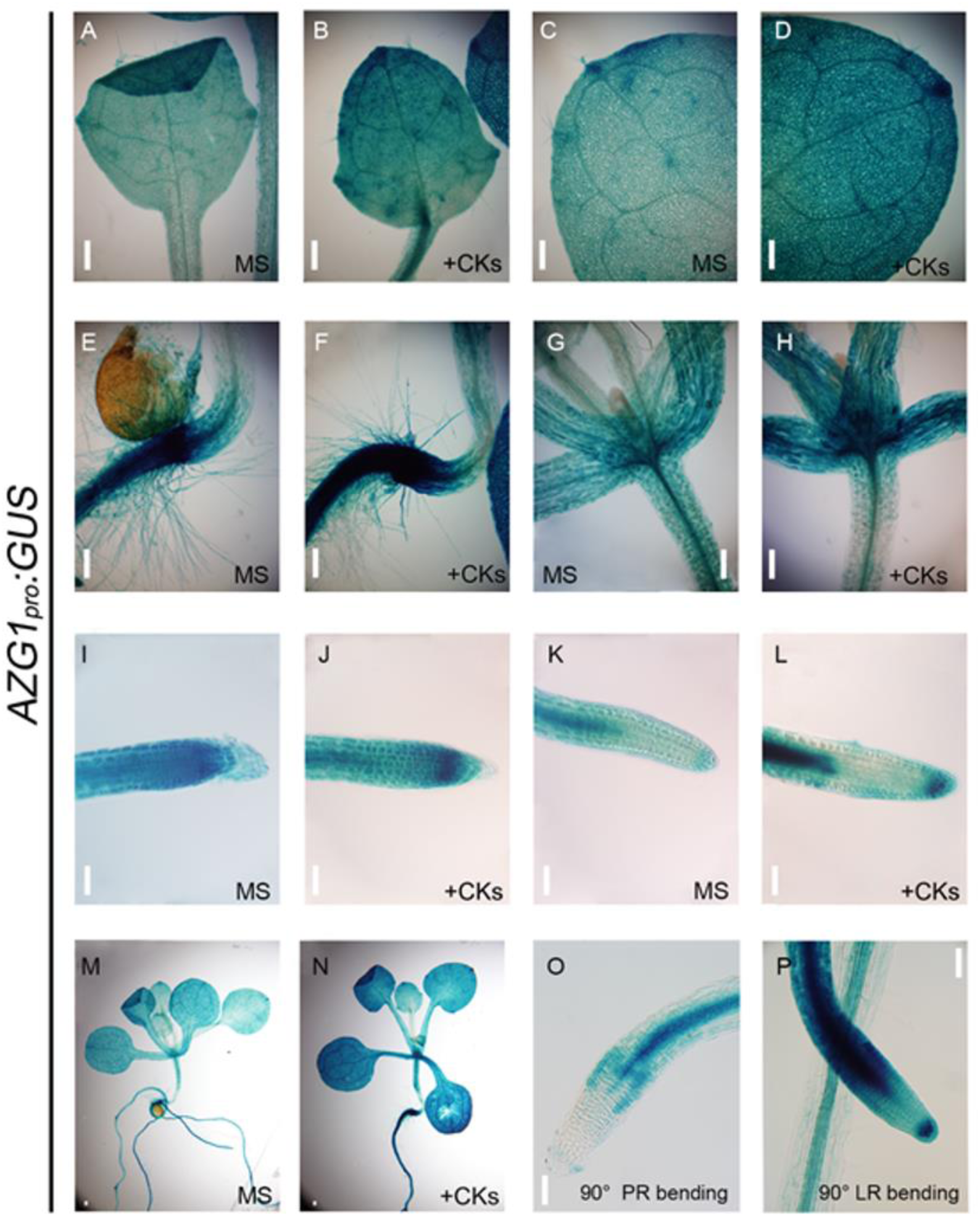
*AZG1pro:GUS* expression in Arabidopsis. Expression of GUS reporter gene under the control of an *AZG1* promoter in (A-D) leaves, (E-F) transition zone, (G-H) hypocotyls and shoot apical meristem, (I-J) primary root apical meristem, (K-L) secondary root apical meristem, (M-N) entire seedling and after a 90° rotation of (O) the primary root (PR) or (P) a lateral root (LR). The legend +CK indicates a 4 h incubation with 200 nM tZ. Scale bar = 100 µm.

### AZG1 stabilises PIN1

During LR organogenesis, cytokinin redirects PIN1 for degradation (Marhavý et al., 2011; Marhavý et al., 2014). Though the mechanism of this regulation is known to involve post-translational processes, its exact nature remains unclear. As a protein which simultaneously binds PIN1 and cytokinin in the plasma membrane, AZG1 is a potential point at which this control could be effected. The genomes of flowering plants encode two AZG homologs, named AZG1 and AZG2. To test whether either protein affected PIN1 stability in Arabidopsis, the relative amount of PIN1 was visualized in three-day-old seedlings of WT, *azg1-1*, and *azg1-1* x *azg2-1* plants. Immunolocalization of PIN1 in *azg1-1* and *azg1-1* x *azg2.1* genotypes indicated a progressive contraction of the PIN1 expression domain (Figure S4), and western blot analysis confirmed that two-day old *azg1-1*, *azg2-1* and *azg1-1* x *azg2-1* seedlings contained less PIN1 than did WT seedlings of the same age (Figure 3L). Overexpression of *AZG1* using the 35S promoter had no observable effect on PIN1 abundance (Figure 3L). Blocking proteasome-mediated protein degradation by treating plants with MG132 for 90 minutes resulted in the reappearance of PIN1 in *azg1-1* x *azg2-1* seedlings (Figure 3M). We therefore conclude that AZG1 and AZG2 act together to stabilize PIN1. Blocking protein synthesis in seedlings by treating them for 90 minutes with cyclohexamide did not visibly affect the amount of PIN1 in WT plants, suggesting that the protein was stable in the membrane over this time period. Over the same period, cyclohexamide treatment had no effect on PIN1 abundance in *azg1-1* or in *azg1-1* x *azg2-1*.

### Cytokinin responsiveness during root growth and cell differentiation

CK exerts a strong and pervasive influence over RSA (Laplaze et al.,2007; Argueso et al., 2009; Werner et al., 2010). In accordance with this, the developing a*zg1-1* root appeared to be particularly sensitive to environmental cues such as the combination of high nitrate (20 mM KNO3) and short photoperiod (8h/16h light/dark), showing a longer primary root and denser LRs when compared to WT (Figure 5A and B). These differences were more noticeable in the presence of exogenous tZ (Figure 5 C-F). *35Spro:AZG1* showed an increased sensitivity to tZ (Figure 5 C-F). Primary root growth inhibition by tZ was restored in a complementation line in the *azg1-2* genetic background (Figure 5F). These results are consistent with endogenous AZG1 increasing the permeability of plasma membranes to CK in developing LR primordia and within the Arabidopsis RAM.

**Figure 5.**
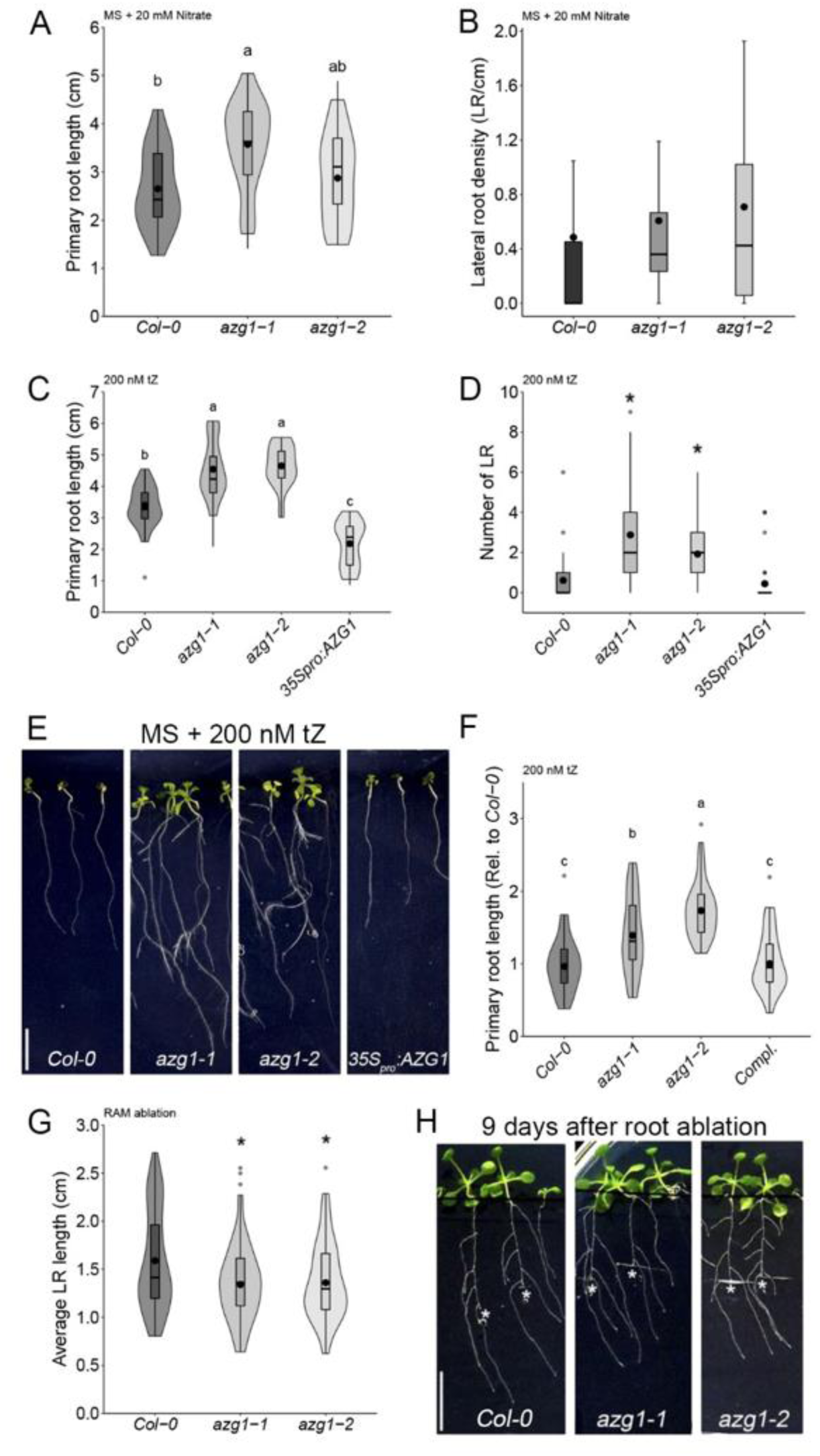
Root phenotype of *azg1* mutant lines. (A) Primary root length and (B) LR density of azg1-1 and azg1-2 growing in 0.5x MS containing 20 mM KNO3 as their sole source of nitrogen. Response of azg1 mutants root architecture without (C) and in the presence of (D and E) exogenous 200 nM tZ treatment with respect to (D) primary root length and (E) number of emerged LR per plant. F) Complementation of primary root length phenotype in the presence of 200 nM tZ (azg1-2 line transformed and selected in homozygotes for AZG1pro:AZG1-GFP. G) Representative plants of Col-0, *azg1-1*, *azg1-2* and *35Spro:AZG1* grown in presence of 200 nM tZ (H) Representative plants after root apical meristem removal. Scale bars represent 1 cm. Asterisk shows significant differences determined by ANOVA test, followed by a Duncan multiple range test (p< 0.05; (A-B) nWt=31, n*azg1-1*=41, nazg1-2=34; (C) nWt=27, n*azg1-1*=28, n*azg1-2*=27, *n35Spro:AZG1*=30; (F) nWt=35, n*azg1-1*=32, n*azg1-2*=27, nCompl.=36) or (D and G) Kruskal-Wallis test, followed by a Wilcox multiple range test ((D) p<0.01; nWt=27, n*azg1-1*=28, n*azg1-2*=27, n*35Spro:AZG1*=30; (G) p<0.05; nWt=54, n*azg1-1*=58, n*azg1-2* =34).

The delicate balance between cell division and cell fate acquisition in is under the influence of diverse signaling molecules (van den Berg et al., 1997; Xu et al., 2006). Among these, the balance of auxin and cytokinin is again prominent in determining cells’ developmental fate, with auxin promoting the formation of roots and cytokinin promoting the formation of shoots (dello Ioio et al., 2008; Murashige and Skoog, 1962). We therefore prepared de-differentiated callus cultures, predicting that *azg1-1* calli would be less sensitive to cytokinin than WT calluses, and therefore more likely to develop roots when treated with kinetin. When treated with media supplemented with kinetin at concentrations of between 230 nM and 3.7 µM, between 50 and 80% of WT root explants grew leaves (Figure S5). In contrast, *azg1* root explants grown on 230 nM kinetin consistently failed to develop aerial tissues. Kinetin at 460 nM was able to stimulate leaf development in 9% of *azg1-1* calli, with a maximum rate of leaf development observed at 930 nM kinetin (Figure S5). We therefore conclude that in WT root explants, AZG1 serves to increase the permeability of the plasma membrane to an inward flux of kinetin.

Observations of the RAM after ablation have previously yielded valuable insights into the way in which relevant signals interact (Xu et al., 2006; Sauer, 2006; Rosquete, 2013). To integrate the dynamics of AZG1 expression and its role in CK signaling into RAM signaling programs, we excised the primary RAM of Arabidopsis seedlings and observed the response of *AZG1pro:GUS* and *TCSnpro:GFP* reporters. After the ablation of the RAM, a decrease in *AZG1pro:GUS* expression was observed both in the main root and in LRs (Figure S6A). Similarly, *TCSnpro:GFP* expression in the stele was down-regulated following RAM ablation, and was barely detectable in non-vascular tissues (Figure S6B). To understand further the causal relationship between changes in the distribution of CK signaling maxima and AZG1 transport activity, we excised the RAM of seven-day old *azg1-1* plants. Nine days later, *azg1-1* plants developed shorter LRs when compared to WT plants (Figure 5G). These data suggest that AZG1 plays a role in the remobilization of CK to sites of lateral root development during root system reorganization after loss of the primary RAM.

### Cytokinin signaling in the RAM and lack of RAM phenotype

To test the hypothesis that AZG1 drives a cytokinin signaling maximum in the RAM, plants expressing the *ARR5pro:GUS* reporter were crossed into both *azg1-1* and *35Spro:AZG1* plants. In all cases, *ARR5*-driven GUS staining was localized to columella cells (Figure 6). The distribution of staining was narrower in *azg1-1* plants when compared to WT, where staining was also observed in stele stem cells (Figure 6, A and D). In contrast, *35Spro:AZG1* plants displayed a broader distribution of GUS-dependent staining, with coloration being observed in stele cells of the meristematic division zone (Figure 6, G). A similar observation was also made in two independent insertion lines of the *TCSnpro:GFP* reporter in the *azg1-1* background. Defective CK signalling has been reported in the stele region in CK-deficient mutants *log3/log4* and *lhw* mutants (Ohashi-Ito et al., 2014). In *azg1-1* lines, CK signalling in stele cells of the root meristematic zone was heavily repressed (Figure S7 A-B).

**Figure 6.**
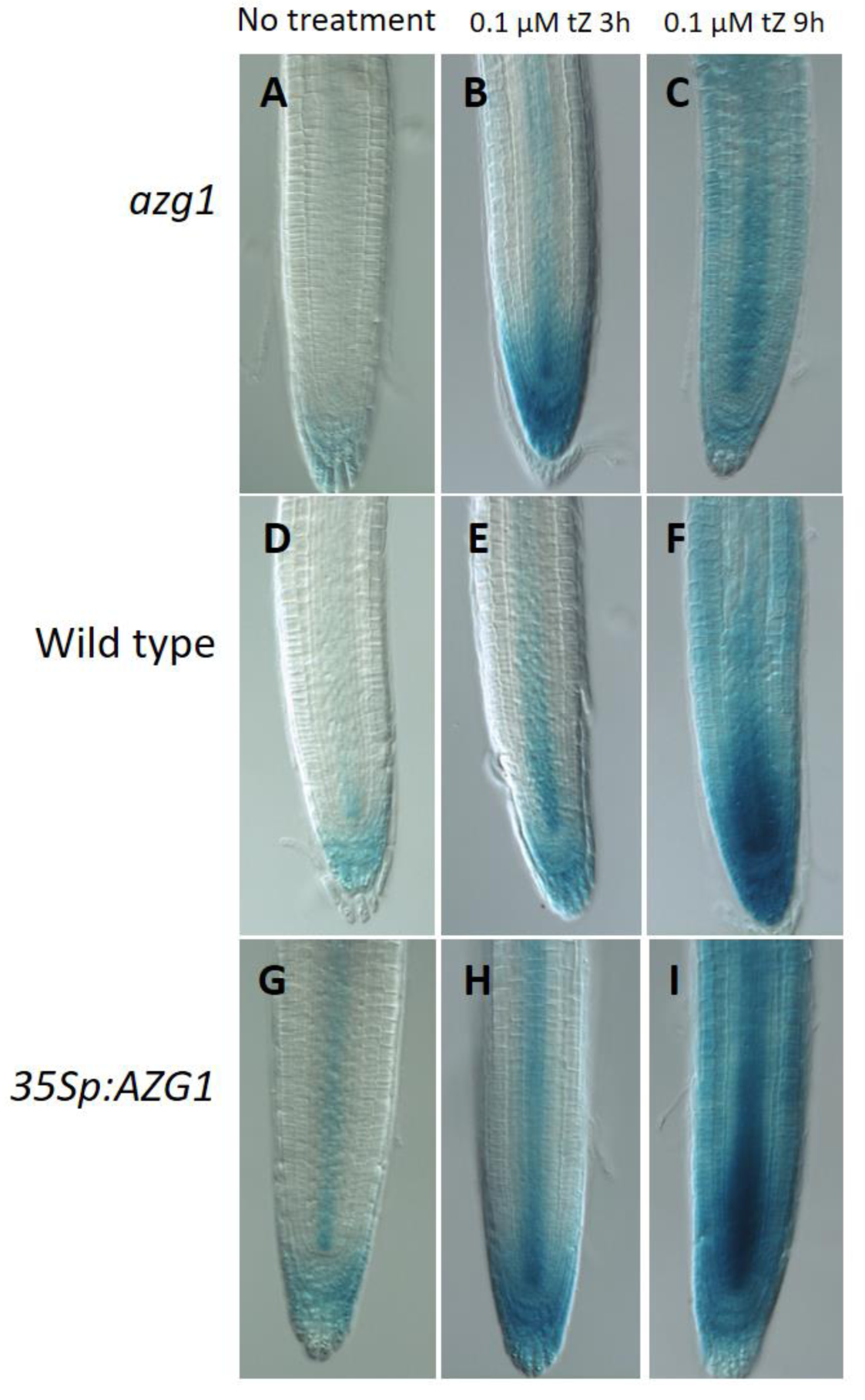
AZG1 drives cytokinin signaling in the RAM. A) Localization in Arabidopsis roots of *ARR5pro:GUS* expression in *azg1-1* (A-C), WT (D-F), or *35Spro:AZG1* (G-I) genetic backgrounds. Roots were either untreated (A, D, and G), or treated with 100 nM tZ for either 3 hours (B, E and H) or 9 hours (C, F and I).

#### ARR5pro

*GUS*-dependent staining in seedlings was cytokinin dependent after treatment with 100 nM tZ for either 3 hours (Figure 6, B, E, H) or 9 hours (Figure 6, C, F, I). In tZ-treated *35Spro:AZG1* plants, the cytokinin-dependent signaling maximum extended through the vascular cells of the division zone and into those of the elongation zone (Figure 6 H and I).

To test the effect on the structure of the RAM of losing AZG1 function, the cellular architecture of populations of *AZG1* and WT roots were compared. To do this, iRoCS, a pipeline which assigns root cells 3D coordinates relative to the root’s central spline, was used. Surprisingly, and despite a difference in the distribution of cytokinin-dependent signaling maxima, no significant difference between the cellular structures of the *azg1-1* and WT RAM was observed (Figure S8).

### AZG1 expression is induced by NaCl

Cytokinin influences plants’ ability to tolerate NaCl in the soil (O’Brien and Benková 2013). We therefore next investigated whether AZG proteins mediated changes in the architecture of the root system, especially those in response to increased salinity or decreased water availability. For this purpose, seven day old WT and *azg1-1 x azg2-1* plants were transferred to ½ MS medium containing either 25 or 50 mM NaCl, and the density of fully emerged LRs was measured after a further ten days of growth. In order to parse the effects of decreased water availability and the presence of NaCl, the experiment was also conducted in the presence of either 50 or 100 mM mannitol.

In the absence of mannitol WT plants, LR density decreased as a function of increasing NaCl concentration, whereas no significant difference was measured in *azg1-1x azg2-1* roots (Figure 7A). The effect of the *azg1-1 x azg2-1* mutations on LR density depended on the salt concentration: in the absence of NaCl, LR density was lower in *azg1-1 x azg2-1* than in WT, but at 50 mM NaCl it was higher (Figure 7A).

**Figure 7.**
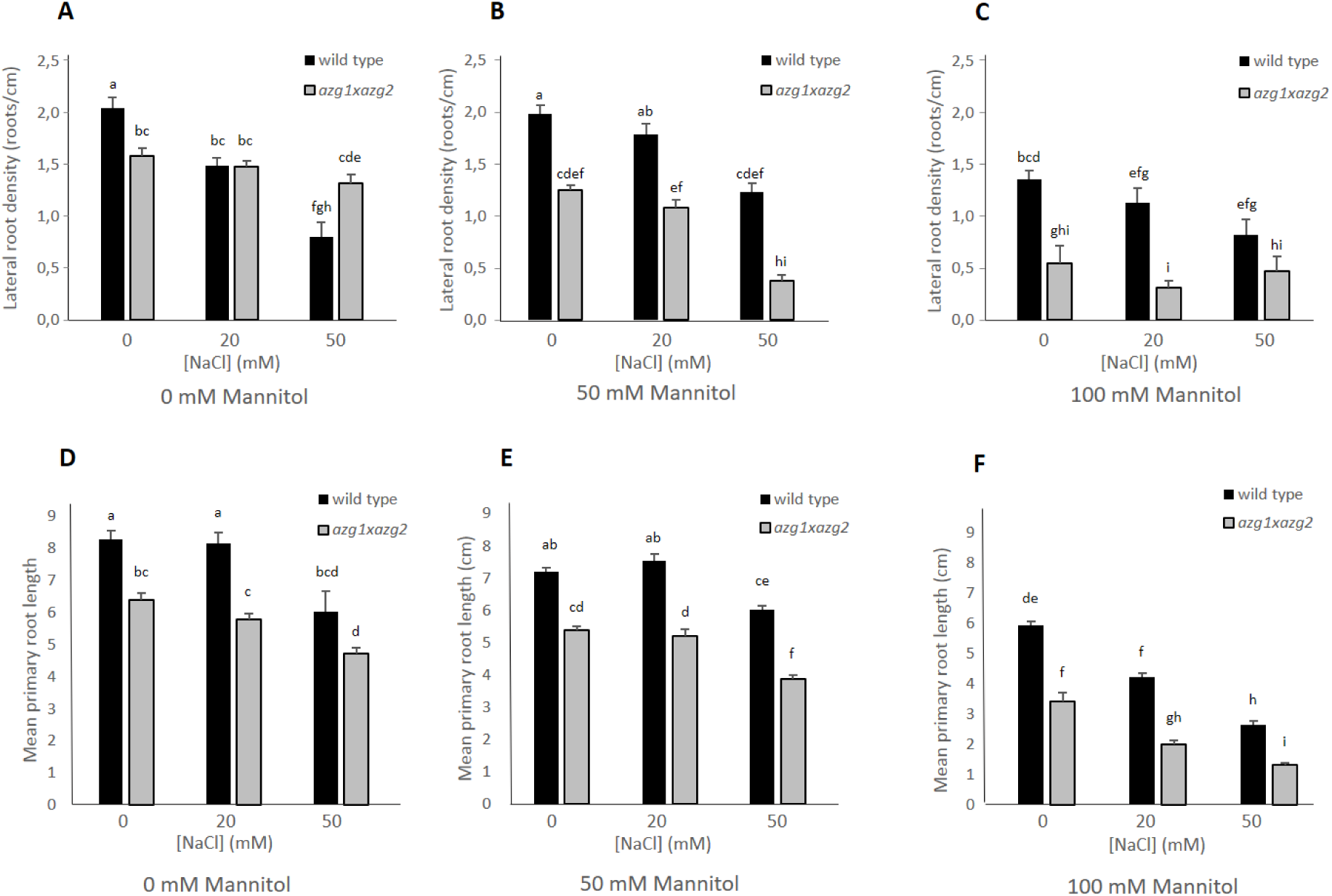
AZG1 couples the response to drought and salinity in Arabidopsis roots. Seven-day old Arabidopsis plants were transferred to solid ½ MS containing combinations of 0 mM, 20 mM or 50 mM NaCl and (A, D) 0 mM mannitol, (B, E) 50 mM mannitol, (C, F) 100 mM mannitol and grown for a further ten days. Letters indicate significantly different densities (*p<*0.01) after a Tukey test. 22<n<58.

Supplementing media with NaCl has two simultaneous effects: those due to sodium ions (sodicity), and those due to a simultaneous reduction in water availability. In order to parse these effects, plants were transferred to plates supplemented with mannitol in order to alter sodicity and water availability independently. In the presence of 50 mM mannitol (a medium with the same osmotic strength as 25 mM NaCl), LR density was inhibited by the addition of NaCl at either 25 mM and 50mM NaCl in WT and *azg1-1 x azg2-1* (Figure 7B). This effect was also observed at 100 mM mannitol, with LR density also being inhibited by NaCl in a similar manner in both WT and *azg1-1 x azg2-1* plants (Figure 7C).

By comparing LR density at similar osmotic pressures, and simultaneously at different NaCl concentrations, (50 mM NaCl (see Figure 7A); 25 mM NaCl + 50 mM mannitol (see Figure 7B); 100 mM mannitol (see Figure 7C)), we were able to compare the effects of sodicity and drought. When osmotic pressure was kept constant by mannitol in this way, the presence of NaCl lowered LR density in WT plants, but increased it in *azg1-1 x azg2-1*. Only the strength but not the direction of the root growth response to mannitol and/or NaCl was affected when WT and *azg1-1 x azg2-1* plants were compared (Figure 7 D-F). We therefore conclude that the ability of WT roots to decrease LR density in response to NaCl requires AZG proteins. This result identifies AZG1 as a potentially important factor in the response of the LR system to local increases in NaCl concentration in the soil. Such an effect was not observed when considering main root length (Figure 7D-F).

In order to test whether *AZG1* gene expression was altered upon exposure to NaCl in the region of the developing LR *AZG1pro*-dependent GUS transcription was observed. Transcription was not observed in the area which surrounded the emerging LR primordium (Figure 8A and B), but became localised to vascular cells at the base of the elongating LR (Figure 8C).

**Figure 8.**
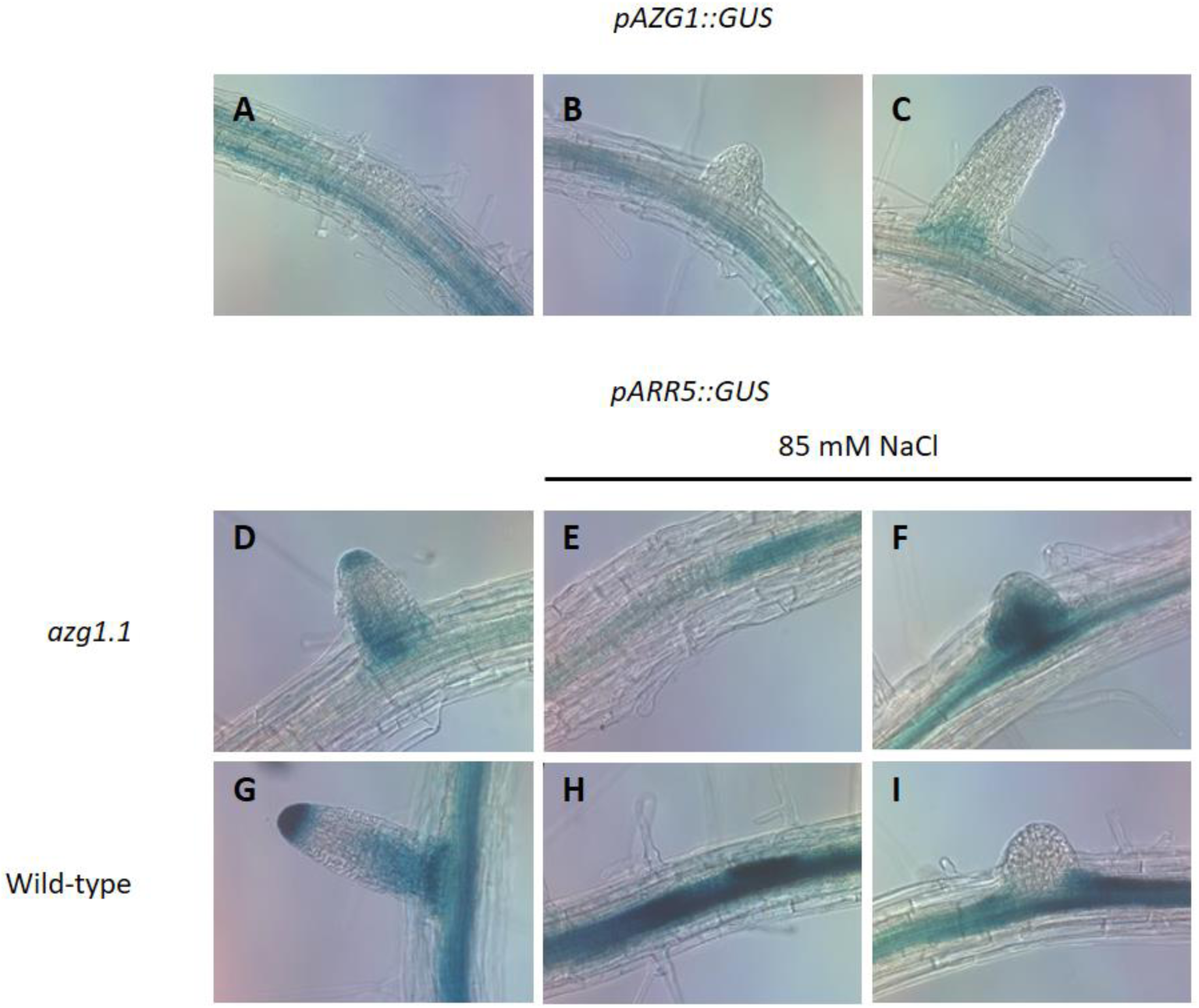
AZG1 *influences lateral signaling in response to environmental sodium chloride*. (A-C) *AZG1pro:GUS* staining in the emerging arabidopsis lateral root. *ARR5:GUS* staining in young lateral roots and their primordia in response to 85 mM NaCl (*aq*). (D-F) WT; (G-I) *azg1-1;* no treatment (D and G); (E, F, H and I) after growth on 85 mM NaCl.

*ARR5*-dependent cytokinin signalling in vascular cells below the developing LR was dependent on the presence of *AZG1* expression (Figure 8D and G). The application of 85 mM NaCl had a conspicuous effect on the distribution of *ARR5pro:GUS*-dependent staining in the emerging LR, with staining in *azg1-1* in the presence of 85 mM NaCl being localised to the emerging LR primordium (but absent from the stage II primordium) and being absent from subtending vascular cells on the proximal side of the root with respect to the emerging LR (Figure 8, E and F). This contrasted markedly with the staining pattern in the WT background, which displayed staining throughout the subtending vasculature and no staining in the emerging LR (but staining in the stage II primordium) (Figure 8, H and I).

## Discussion

### Cytokinin transport

Two principal long distance cytokinin (CK) transport streams have been characterized in flowering plants: those of *trans*-zeatin (*t*Z) from the roots to the shoot via the xylem, and of N^6^(Δ^2^-isopentyl)adenine (iP) and *cis*-zeatin (*c*Z) from the shoots to the roots via the phloem. This transport is primed by members of three families of transporter protein: the purine permeases (PUPs), the equilibrative nucleotide transporters (ENTs), and the ATP-binding cassette (ABC) transporters (Liu et al., 2019). The fact that these streams are both spatially and functionally distinct implies a degree of specificity among transporters for different cytokinin substrates, although quantitative data on transport rates and flux capacity of each transporter is currently sparse.

Despite CKs being synthesized in leaves (Kuroha et al., 2009), a series of careful grafting experiments has shown that the majority of tZ which is present in the shoot is synthesized in roots and transported via the xylem (Matsumoto-Kitano et al., 2008). This work has been corroborated by the quantification of CK in vascular exudates which revealed that iP is the major CK in phloem streams, whereas 90% of an Arabidopsis plant’s *t*Z is present in the in the xylem, where *t*Z-riboside (*t*ZR) accounted for 80% of all CK species measured (Hirose et al., 2008; Kudo et al., 2010; Ko et al., 2014). There is currently no strong evidence available that ENTs are involved in cellular *t*Z uptake (Liu et al., 2019); moreover, those PUPs which have been classified as CK transporters (PUPs1 and 2), are likely to be responsible for the vascular loading of CK in leaves (Bürkle et al., 2003). Although ABCG14 is required for CK-dependent processes in the shoot, its localization supports the hypothesis that it is necessary for the loading of *t*Z into xylem vessels in the elongation zone of the RAM (Zhang et al., 2014). Plants of *abcg14*, which accumulate *t*Z in roots and dissipate *t*Z in shoots, show CK-deficient shoot phenotypes such as thin stems containing a reduced number of vascular bundles, small rosettes and small flowers. In contrast to this, here we found that *azg1* loss of function plants do not show such differences. These physiological observations, along with reduced *t*Z accumulation rates and resistance of root growth to exogenously applied tZ in *azg1* lead us to conclude that AZG1 is responsible for the cellular import of *t*Z in the RAM, whereas ABCG14 is responsible for its cellular efflux. The expression domains of AZG1 and ABCG14 also imply their involvement in different processes: while AZG1 is localized to procambium cells in the division zone of the RAM (Figure 2D), ABCG14 is localized further from the root tip in cells of the elongation zone (Zhang et al., 2014). The expression pattern of AZG1 within the RAM largely resembles that of the CK signaling reporter *TCSnpro:GFP* (Figure S7; Zürchner et al., 2013). *t*Z is supplied to the shoots from the roots (Beck and Wagner, 1994; Beveridge et al., 2000). Our data do not support the involvement of AZG1 in the long-distance shootward transport of this *t*Z; but rather a role for AZG1 in the retention of *t*Z in via its cellular reuptake in the RAM (Figure 9A).

**Figure 9.**
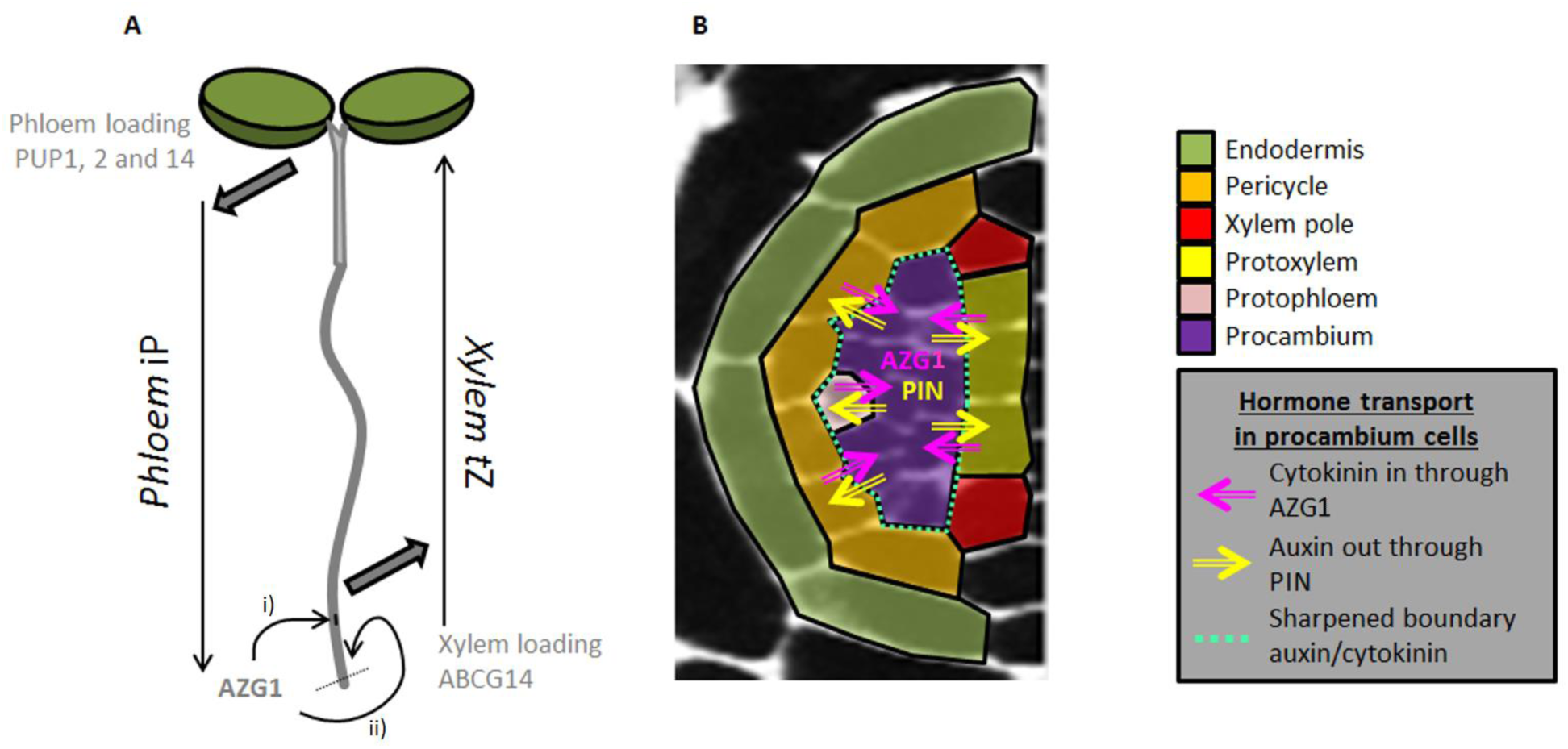
The context of AZG1-dependent cytokinin transport. A) Cytokinins are actively loaded into both phloem and xylem streams for bulk long-distance transport. The cellular localization of AZG1 suggests it is not involved in these processes, but rather in maintaining cytokinin signalling maxima in i) developing lateral roots and ii) the root apical meristem. B) AZG1 is localized to procambium cells where it acts simultaneously to stabilize cellular auxin efflux and cytokinin influx. This could affect sharpened boundary between auxin maxima in developing xylem cells, and cytokinin maxima in developing phloem cells.

### Cytokinin and patterning of the RAM

This hypothesis may, however, easily be challenged as our careful cell-by-cell analysis showed that *azg1* plants show no distinctive phenotype within the RAM. CK affects many cellular processes in the root including cell division, LR development, the response to abiotic stresses, and nodule development (Kieber and Schaller, 2018; Sasaki et al., 2014). It also shortens the length of the meristem itself in a process which is mediated by the transcriptional regulation of *SHY2*, a repressor of auxin-dependent transcription (dello Ioio et al., 2008). Surprisingly, and in contrast to the case seen in *abcg14*, the RAM of *azg1* displayed a more constrained domain of CK signaling (as visualized by *pARR5:GUS and TCSnpro:GFP* expression), which one might have expected to confer an increased meristem size. That it did not, could be interpreted in terms of the functional significance of the AZG1-PIN1 interaction. AZG1 stabilizes PIN1 by reducing its rate of proteasome-dependent degradation. This observation would be consistent with a decrease in cellular auxin efflux in *azg1* plants; a hypothesis consistent with our observations would therefore involve the PIN1-AZG1 interaction buffering cellular auxin:CK ratios via the coupling of auxin efflux regulation and CK influx within the RAM. The presence of additional CK transporter(s) at the plasma membrane during embryogenesis goes some way to rationalize the correlation which was observed between the localization of PUP14 in heart-stage embryos and areas of low CK signaling with the increasingly uncontroversial hypothesis that CK perception is localized at the ER (Zürchner et al., 2016; Romanov et al., 2018).

### Posttranslational control of PIN1 by cytokinin

The bilateral symmetry of vascular cells within the RAM is established and maintained by an intricate pattern of cross-talk between auxin and CK-mediated signaling systems (Vaughan-Hirsch et al., 2018). At its core is the spatial separation of auxin and CK signaling cascades; auxin-mediated signals are localized to protoxylem cells, and CK signals to procambial and protophloem cells. This separation is maintained by transcriptionally mediated, but spatially defined, signaling mechanisms such as the induction of AHP6, a repressor of CK signaling, in protoxylem cells (Mähönen et al., 2006; Bishopp et al., 2011a). As well as its activity being thus suppressed in protoxylem cells, CK biosynthesis is up-regulated there by the production of LONELY GUY4 (LOG4), which is stimulated by the auxin-dependent transcription of TARGET OF MONOPTEROS5 (TMO5) and its dimerization with LONESOME HIGHWAY (LHW). In this model, CK then diffuses from protoxylem into procambial cells, where it is free to stimulate transcription.

AZG1 is localized to the plasma membrane of procambial, but not protoxylem, cells. It may not therefore be simple diffusion which mediates the crucial step of CK transfer between cell types, but an active uptake which is specifically confined to procambial cells. The binding and stabilization of PINs by AZG1 may also play a role in a second aspect of the proposed model. PIN7 relocalises in procambial cells in order to focus auxin into protoxylem cells (Bishopp et al, 2011b). The post-translational stabilization of PINs and AZG1 in procambial cell into lateral domains of mutual auxin efflux and CK influx would certainly be consistent with the proposed model (Figure 9B).

That CK influences the post-translational regulation of PIN proteins is already established. In stage III LR primordia, CK stimulates the removal of PIN1 from anticlinal membranes (Marharvý et al., 2014). This phenomenon is not cell-type specific, but is highly dependent on the sub-cellular localization of PIN proteins. Going forward it will be a priority to test whether the AZG1 dependent post-translational stabilization of PIN1 directly promotes LR emergence, and whether either subcellular AZG1 localization or AZG1-mediated cytokinin influx can mediate the post-translational response of PIN1 to cytokinin.

Despite displaying a restricted procambial cell CK signaling domain, and unlike other CK-related mutants with a similar restriction in their CK signaling domain (*e.g. log3*/*log4*/*log7*, *lhw*; Ohashi-Ito et al., 2014), *azg1* seedlings show no RAM phenotype. As AZG1 potentially regulates auxin and CK transport and signaling processes simultaneously, it is conceivable that auxin: CK ratios may be only minimally affected in crucial cell types in *azg1* when compared to WT roots. This simultaneous regulation may contribute to the robustness of root development. The localization of AZG1 in procambium cells of the RAM, together with a lack of any clear aerial phenotype, is probably more consistent with a role in the stabilization of tissue patterning mechanisms than in long distance CK transport. However, the possibility that AZG1 is involved in phloem unloading of iP remains open. Finally, a hypothesis involving the redundant effect of diverse CK transporters, allowing a lower but sufficient rate of CK influx, cannot yet be discarded.

### Salt, cytokinin and root system architecture

Cytokinin signaling has previously been broadly linked to plant responsiveness to the environment (Cortleven et al., 2019), as external conditions modulate CK homeostasis, for instance by altering its biosynthesis (Takei et al., 2004, Kuppu et al., 2013). The role of CK transport, and hence its physiological impact, may therefore influenced by environmental cues through the regulation of endogenous CK availability. This influence became evident after two independent investigations into the ABCG14 CK transporter. In these approaches, contrasting root elongation phenotypes were observed in identical *abcg14* loss-of-function genotypes (Ko et al., 20014; Zhang et al., 2014). These differences could be ascribed to the different growth conditions used (e.g. in media composition and light/dark regime), indicating a strong buffering capacity of CK transporters with respect to changes in environmental conditions. Such a capacity may also be inferred for AZG1. When grown under high nitrate but short photoperiod (low C:N balance), *azg1* plants developed longer and more complex root systems when compared to the wt (Fig 5 a and b). This phenotype contrasted with that which was observed when plants were grown in the presence of 0.5 MS media containing sucrose under a long-day photoperiod (high C:N balance; Fig 7A). Under a low C:N conditions, the systemic inhibition of root branching is triggered, increasing the amount of endogenous CK (Zhang et al., 1999; Takei et al., 2004). Under these conditions, lines which display a disturbed response to CK, such as *azg1*, would respond aberrantly to this systemic regulation. This effect has been also described for *azg2*, also recently characterized as being deficient in CK transport; this genotype showed a differential phenotype under a low C:N regime after the application of exogenous CK (Tessi et al., 2020).

Publicly available databases of transcriptional data (e.g. www.bar.utoronto.ca) indicate that AZG1 transcription is up-regulated after one hour of exposure to a 150 mM NaCl solution. These data were borne out by the experiments presented in this report, which recorded changes in the distribution of CK signaling maxima between *azg1* and WT plants, but only after exposure to NaCl. In Arabidopsis, CK plays an important role in the regulation of sodium ion concentration in root cells (Tran *et al*., 2007). However, the relationship between CK and soil salinity is multi-faceted and complex. For example, ARR1 and ARR12, two type-B CK response regulators, increase the accumulation of sodium ions in root xylem cells by repressing transcription of the gene encoding the sodium ion transporter AHK1.1 (Mason et al., 2010). Arabidopsis accessions showing relatively high *HKT* expression also bore fewer LRs (Julkowska *et al*., 2017). The absence of a functional *AZG1* gene shifted CK signalling maxima into emerged LR primordia of salt-treated plants, but away from unemerged primordia. In combination with a loss-of-function *azg2* allele, this conferred a LR system which, though slightly less dense than was observed in WT plants, was largely resistant to external NaCl solutions of up to 50 mM (Figure. 7A). This insensitivity seemed decoupled from drought, as in the absence of NaCl, WT and *azg1-1 x azg2-1* roots responded in the same way when exposed to mannitol concentrations of up to 100 mM (Figure 7A-C). The hypothesis that AZG1 is also involved in CK-based root-shoot communication in response to increased soil sodicity is currently doubtful, as its localization makes it unlikely to be involved in the loading of *t*Z into the xylem. Several morphological mechanisms are based on the well-defined and complementary distribution of auxin and CK signalling domains; however, it is still not known how CKs accumulate in specific cells (Bishopp et al., 2011a; Bishopp et al., 2011b; Bielach et al., 2012; De Rybel et al. 2014; Ohashi-Ito et al., 2014; Chang et al., 2015). Our observations involving salt treatment are consistent with AZG1 playing a role in the establishment and maintenance of sharp auxin-cytokinin concentration boundaries within the developing root through its interaction with PIN1 (Figure 9B).

## Acknowledgements

The authors would like to thank K. Kratzat, Z. Kazimierzak and J. Dürr for valuable and expert technical assistance. This work was supported by Bundesministerium für Bildung und Forschung (BMBF SYSBRA, SYSTEC, Microsystems), the Excellence Initiative of the German Federal and State Governments (EXC 294), the German Research Foundation (DFG SFB746 and INST 39/839,840,841), the National Fund of Science and Technology (FONCyT, Argentina, PICT-2009-0114) and a Grant of the Secretary of Science and Technology of the National University of Córdoba (SECyT-UNC). The work of VGM was supported by the Heisenberg Program of the German Research Foundation (DFG) through grants MA2379/9-1 and MA2379/9-2. The authors declare that no competing interests exist.

## Author contributions

V.M., M.D. K.P. and W.T. conceived the study, V.M., M.D., V.M., T.T. and W.T. wrote the manuscript. T.T., M.S., V.M., E.M., T.P., B.S., K.K., JD., N.F., M.N., Z.K., A.W., M.D. and W.T. Planned and performed experiments and analysed data, T.T., M.S., V.M, M.D. and W.T were responsible for the resources. O.N. and M.S planned and performed metabolomic analysis, JT: planned and performed proteomic analysis, V.M, K.P., M.D. and W.T supervised the study. V.M., K.P. and M.D. acquired the funding.

## Supplementary Figure Legends

**Supplementary Figure 1** Candidate PIN1 interactors which did not show direct physical interaction with PIN1 in co-immunoprecipitation assay. A) ADL1:YFP. Immunoprecipitate of ADL1:YFP was analyzed for the presence of PIN1 by western blotting (WB) with anti-PIN1 antibodies. Asterisk indicates full-length PIN1 protein. B) SMT2:HA. Immunoprecipitate of PIN1:YFP was analyzed for the presence of SMT2:HA by WB with anti-HA antibodies. Asterisk indicates SMT2:HA. Protein extract from untransformed tobacco leaves served as a negative control for immunodetection of HA-tagged fusion protein (WB control).

**Supplementary Figure 2** Relative endogenous cytokinin (CK) content in rosette leaves of WT and *azg1-1* plants grown on solid MS either (A) without or (B) containing 200 nM tZ. Bars show standard deviation. N=4. tZ, *trans*-Zeatin; tZR, *trans*-Zeatin riboside; tZ9G, *trans*-Zeatin-9-Glucoside; tZOG, O-Glucosyl-*trans*-Zeatin; tZROG, O-Glucsyl-*trans*-Zeatinriboside; iPR, Isopentenyl-adeninriboside; iPR-5′MP, Isopentenyl-adeninriboside-5′Monophosphate; iP9G, Isopentenyl-adenine-9-Glucoside. For every cytokinin and derivative, relative values were calculated in comparison to tissue from plants not grown on tZ. Values for WT plants were normalized to a value of 1. Actual values are given in brackets.

**Supplementary Figure 3** A) RT-PCR analysis of AZG1 expression in different plant organs. (B) Bending angle after a 90° rotation of growing plates.

**Supplementary Figure 4** Immunolocalization of PIN1, showing its localization in roots of (A) WT, (B) *azg1-1*, and (C) *azg1-1 x azg2-1* four-day old seedlings.

**Supplementary Figure 5** Leaf regeneration is inhibited in *azg1* calli A) Graph shows *Arabidopsis thaliana* root explants grown on solid callus-inducing medium and transferred to solid ½ MS medium containing the indicated concentration of kinetin and incubated for 18 days. B) An example image of regenerating calli developing leaves in AM supplemented with 0.46 µM kinetin.

**Supplementary Figure 6** *AZG1*-, auxin- and CK-dependent expression in response to root apical meristem ablation. A) *AZG1pro:GUS* B) *TCSpro:GFP* in regenerating root apices. Scale bars = 200 µm.

**Supplementary Figure 7** CK signalling in the *azg1* RAM. (A) Col-0 and *azg1* genetic backgrounds were transformed with the *TCSnpro:GFP* CK reporter. Representative RAMs are shown in (A). White arrows indicate where a change in signalling pattern is observed in meristematic stele cells of *azg1-1*. (B) GFP signal intensity average across the zone highlighted in a dashed yellow rectangle in (A) of WT and *azg1-1* backgrounds. Bar = 100 µm.

**Supplementary Figure 8** iRoCS analysis of root dimensions in the Arabidopsis root apical meristem. (A) Average meristematic cell length, (B) cell-file-dependent meristem length (distance from QC), and (C) cell length as a function of distance from the QC. WT (clear); *azg1-1* (shaded). Bars indicate standard deviation.

