## Supplementary figures and images for "The auxin transporter PIN1 and the cytokinin transporter AZG1 interact to regulate the root stress response"

### Supplemental figures

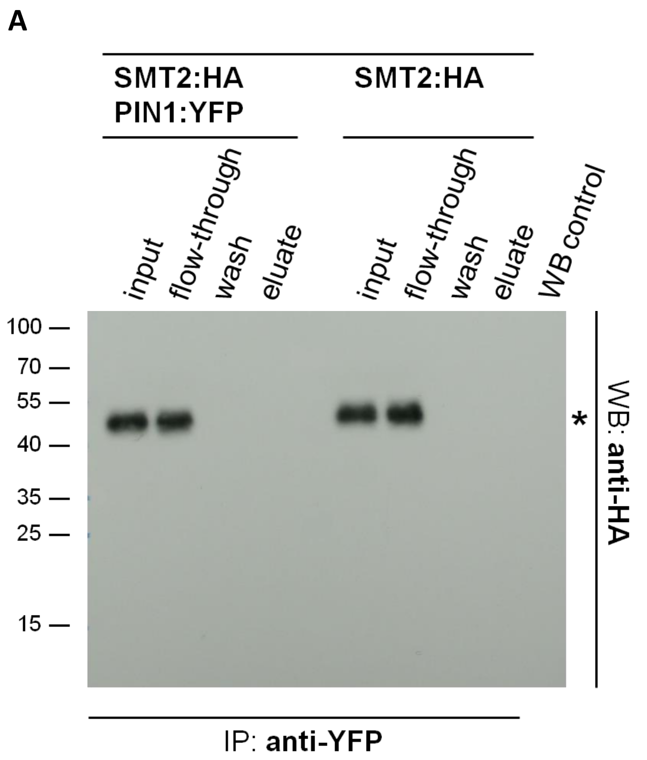

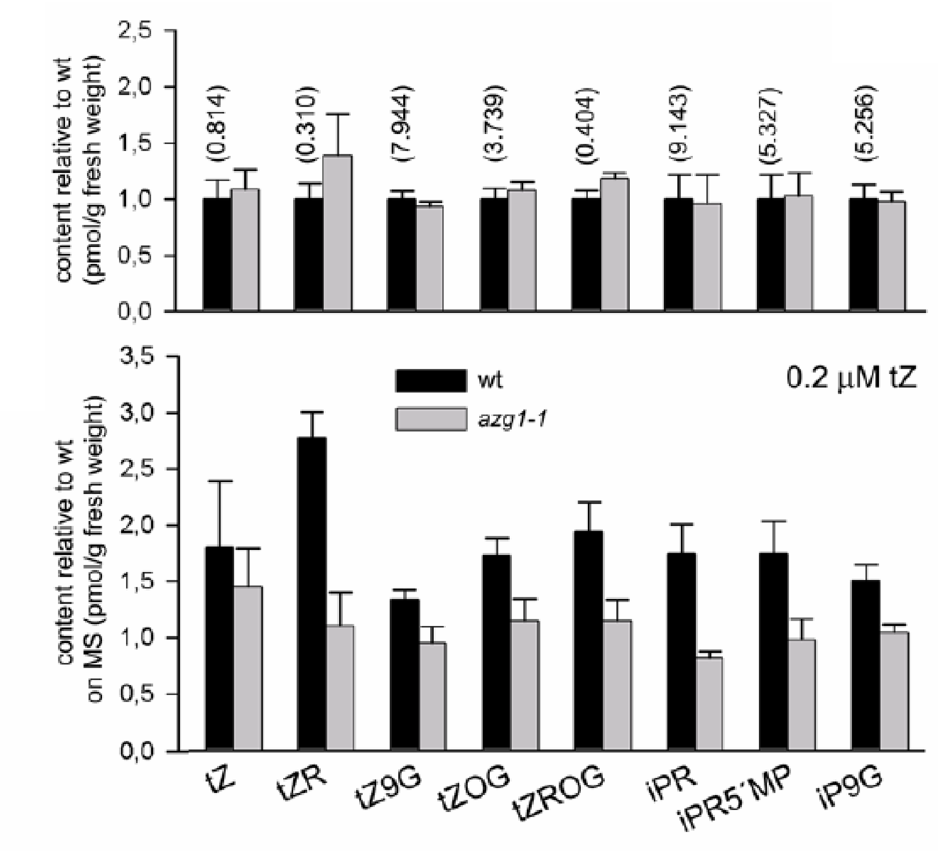

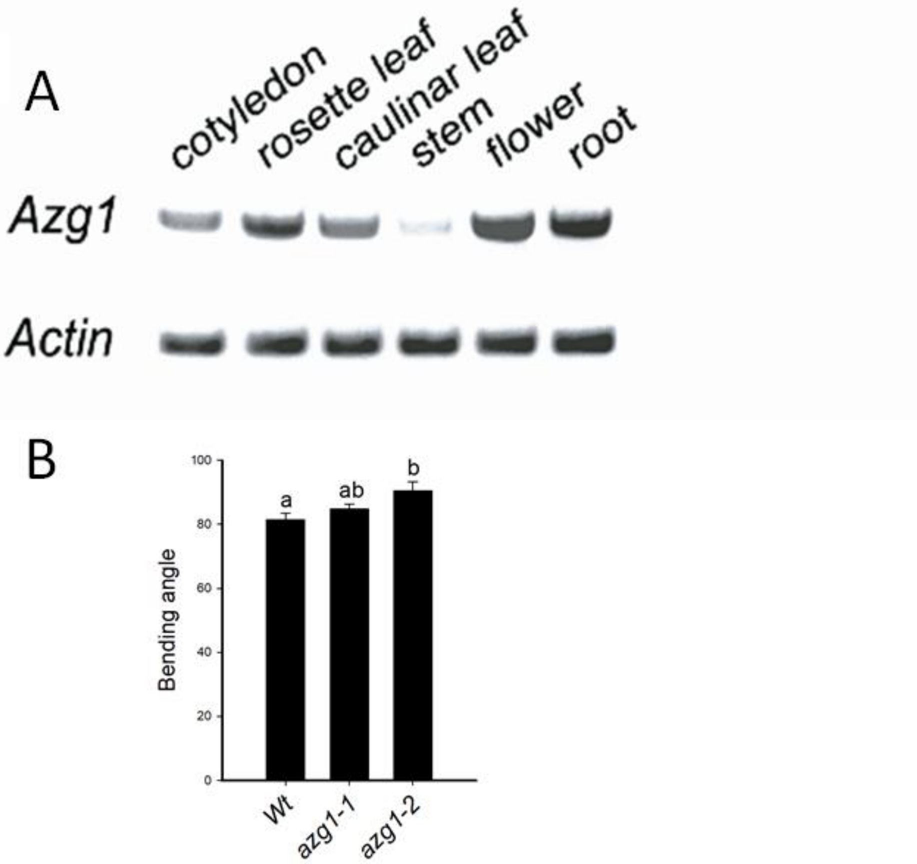

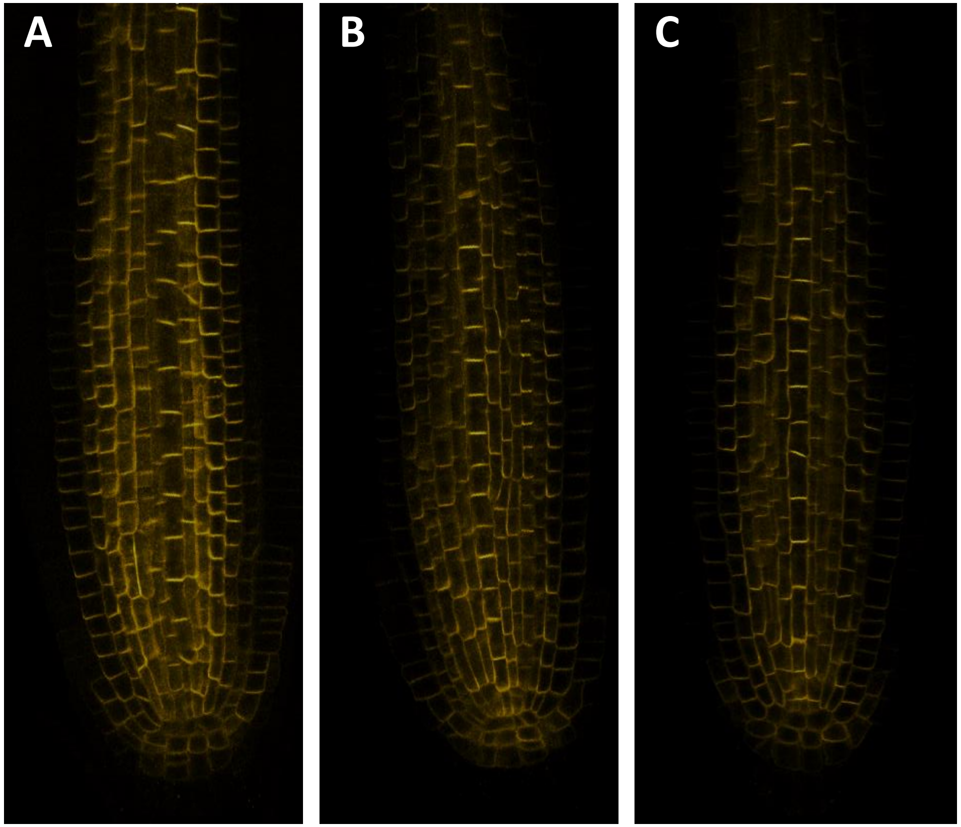

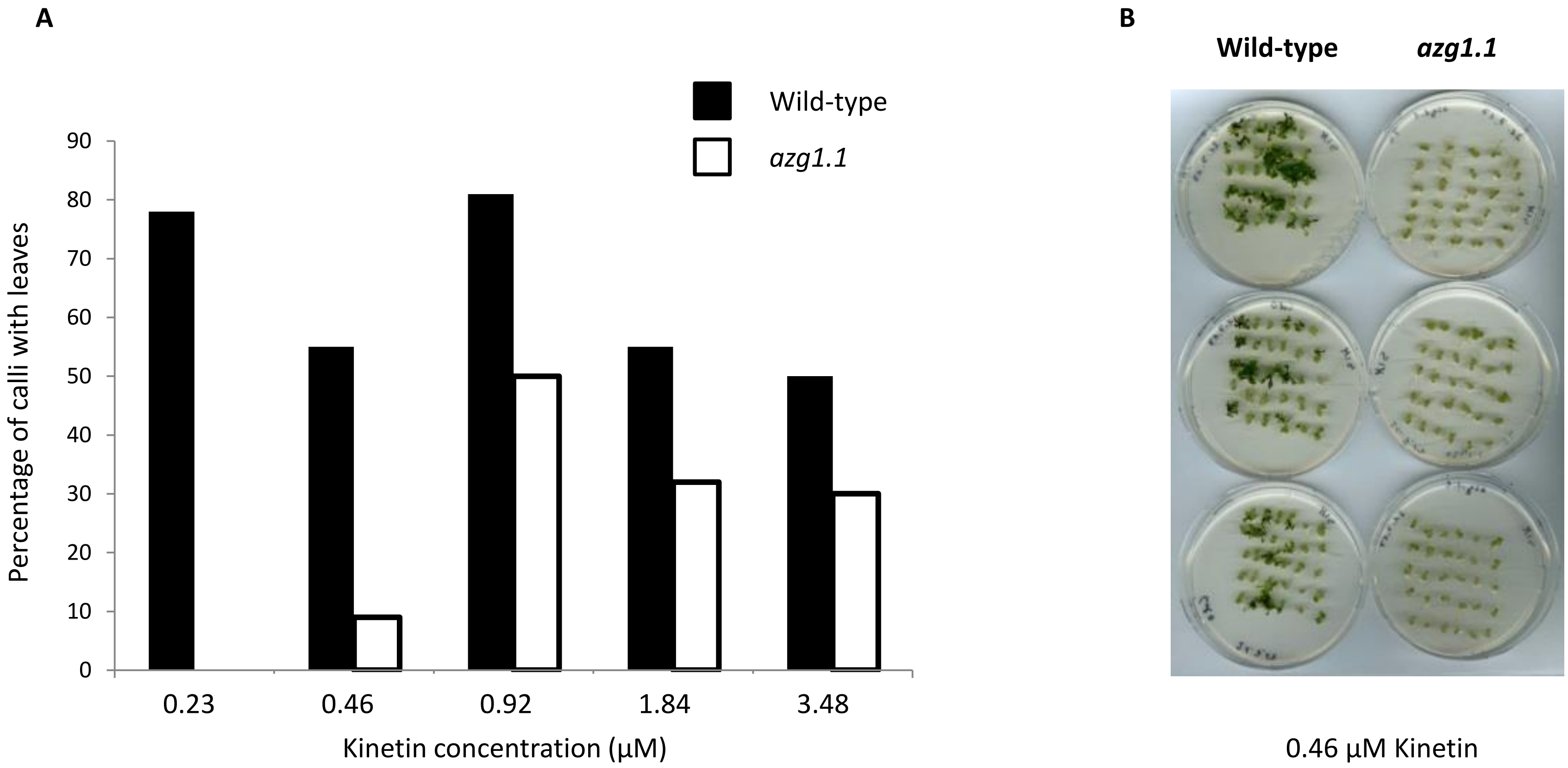

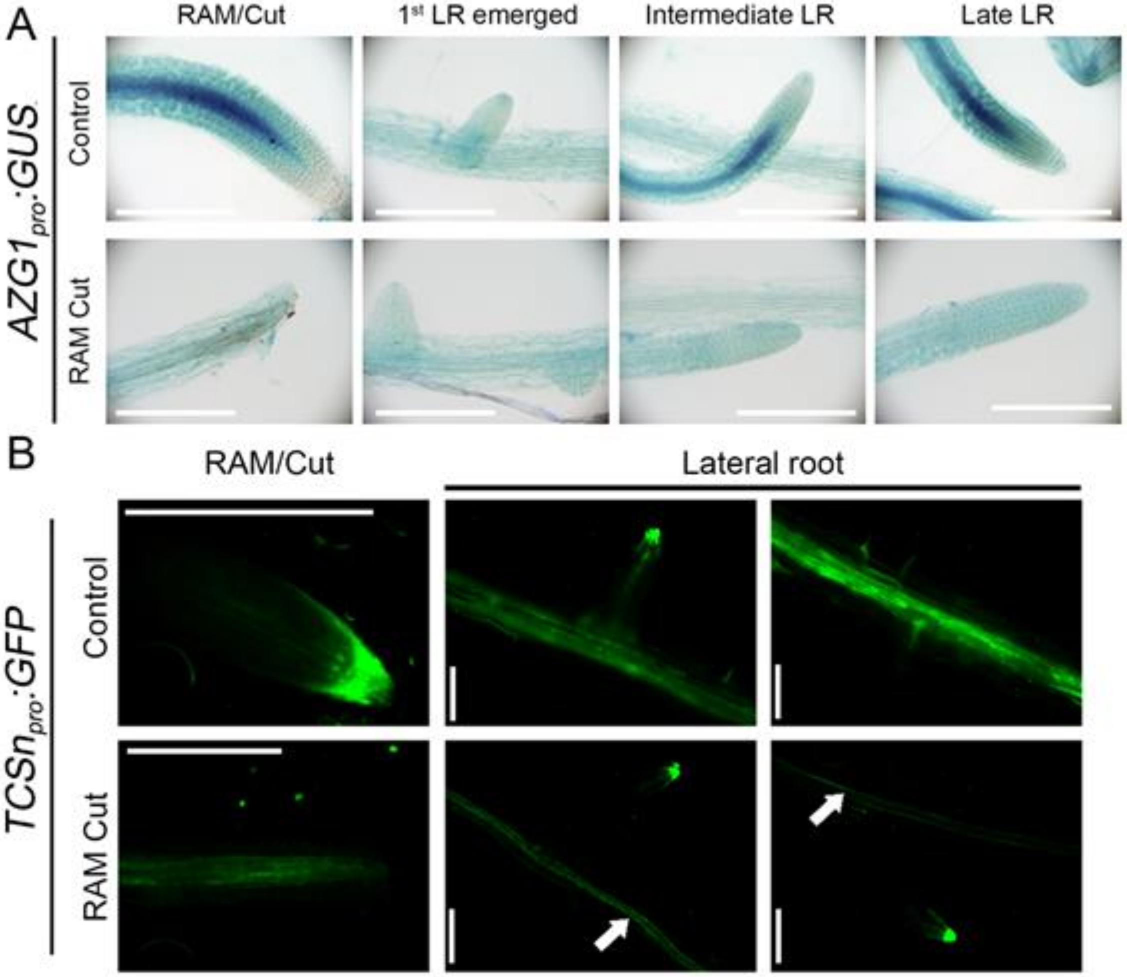

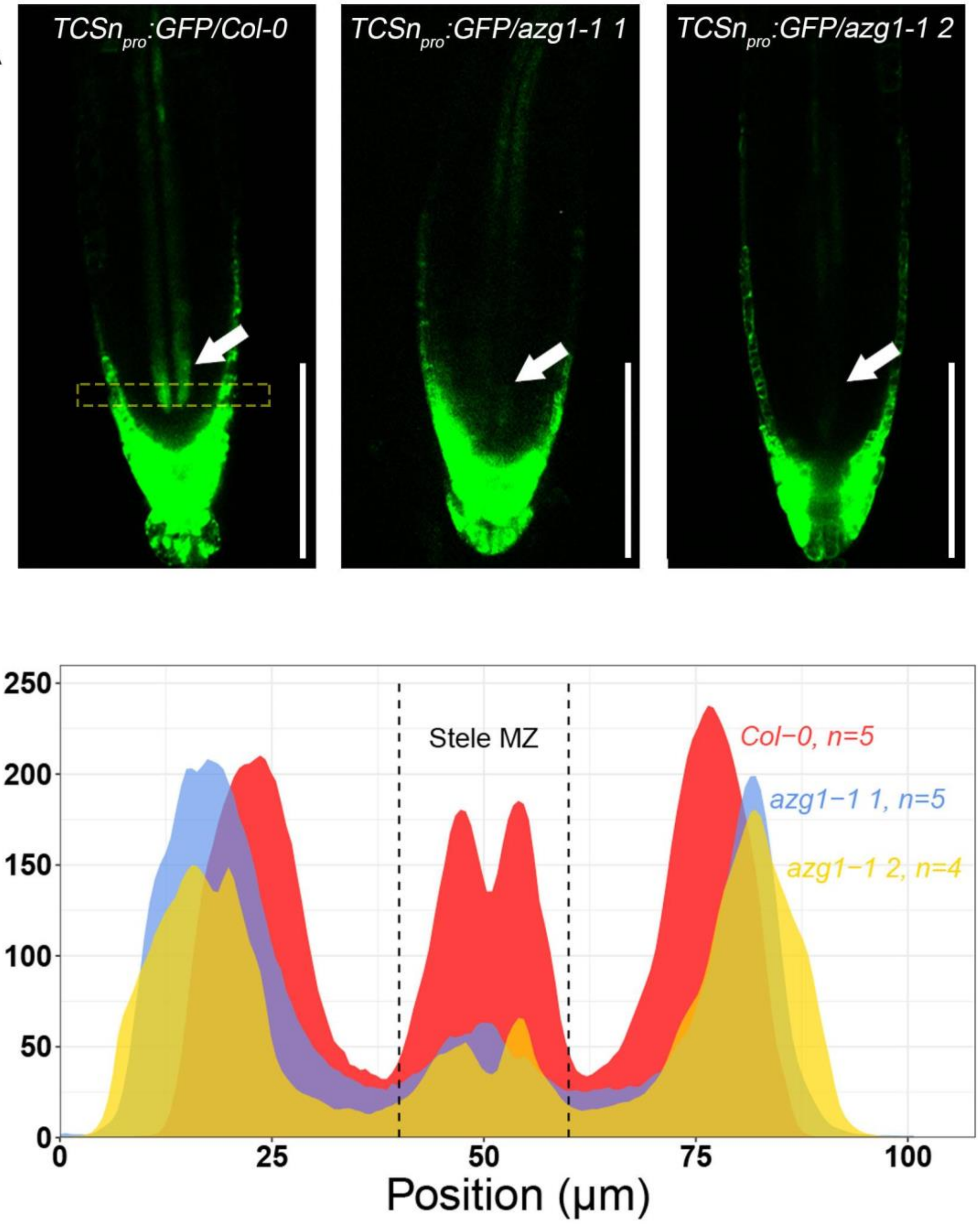

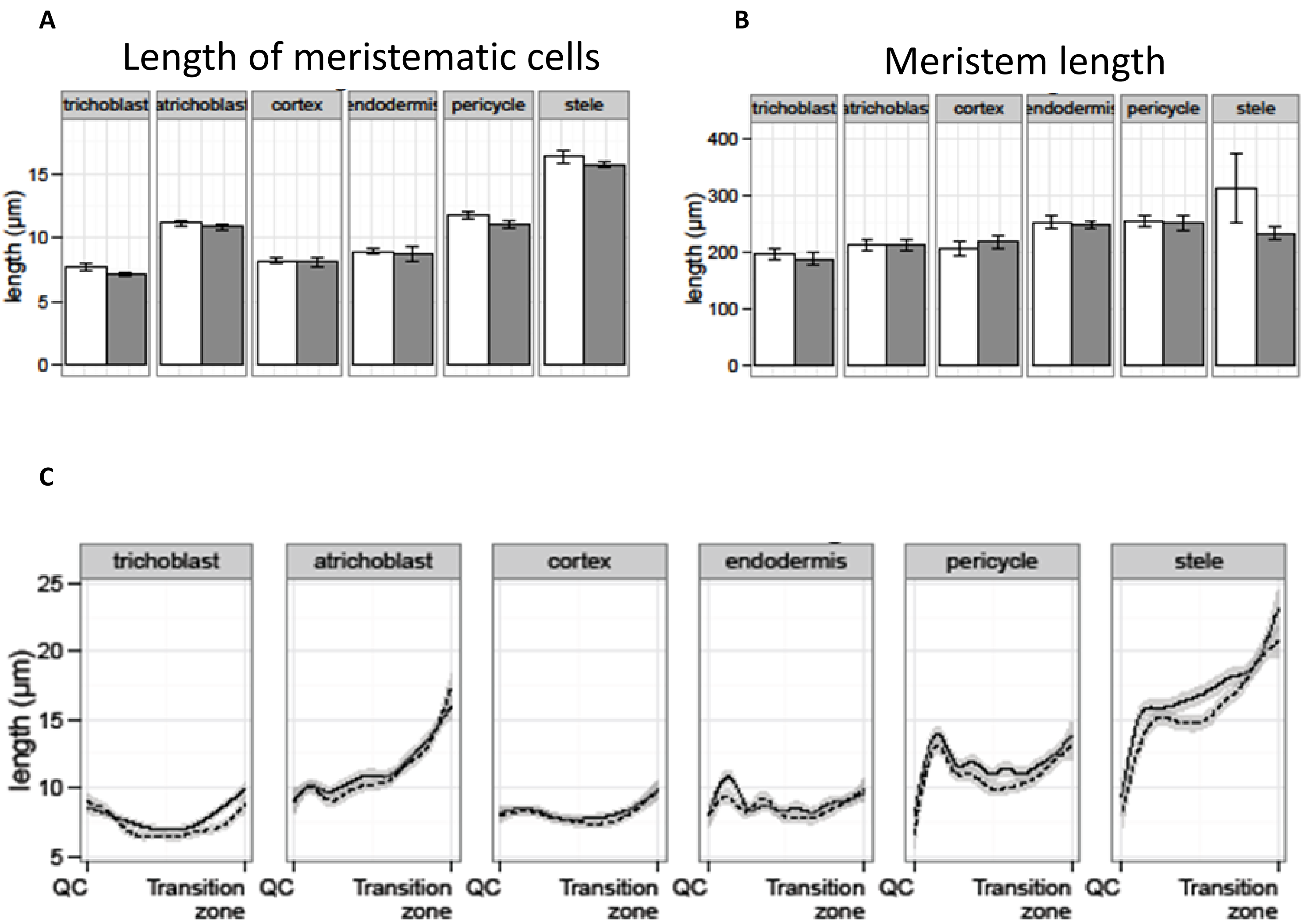
